# Aberrant neuron-OPC synaptic transmission and deficient myelination define sex bias in major depressive disorder

**DOI:** 10.64898/2026.06.14.732204

**Authors:** Cong Zeng, Yinong Li, Xiaorui Wang, Zhonghao Wu, Yan Shen, Susu Li, Shouyu Wang, Xiaoying Chen, Liting Wang, Ying Wan, Alexei Verkhratsky, Jianqin Niu

## Abstract

Sex bias is a notable feature of major depressive disorder (MDD), yet its cellular and molecular origins remain unclear. We integrated computational analyses of human patient data with histological and biochemical validation in animal models to uncover female-specific MDD pathways. We identified female-specific mechanisms involving oligodendroglial lineage cells, especially oligodendrocyte precursor cells (OPCs), as key signal receivers with unidirectional preference in interactome networks. The neurexin (NRXN) pathway emerged as the most perturbed in MDD, with greater disruption in females. Subclustering further identified committed OPCs (cOPCs) as a disease-associated oligodendroglial subpopulation characterized by high expression of synaptic genes; cOPCs were also enriched of MDD-related transcriptional signatures revealed by trajectory analysis. High-dimensional weighted gene co-expression network analysis identified *GABRG3* as an MDD-specific hub gene in cOPCs. The *GABRG3* was also involved in the NRXN synaptic assembly cascade. Using *Gabrg3* conditional knockout mice, we validated findings in patients and demonstrated that conditional deletion of *Gabrg3* in cOPCs recapitulates MDD-like phenotypes, highlighting impaired neuron-cOPC synaptic communication, abnormal myelination, and depression-like behaviors. Together, our work defines a sex-specific cellular and molecular pathobiology of MDD, bridges clinical discovery with preclinical context, and provides a translatable framework for precision medicine targeting fundamental sex differences.

## Introduction

Major depressive disorder (MDD), a main cause of disability worldwide, is a prevalent and debilitating condition characterized by mood disturbances, anhedonia, disrupted physiological functions, cognition, and psychomotor activity ^1^. A notable feature of MDD is the sex-bias, with women being affected at twice the rate of men ^2^; moreover women present greater symptom severity, specific symptomatology, and subjective distress ^3^. Women are also two times more likely than men to experience a single depressive episode and four times more likely to suffer from recurrent MDD ^3^. Despite this well-documented disparity, many depression studies have either excluded female subjects or overlooked sex differences ^4^. The limited data highlight sex-specific differences in various aspects, including genetic architecture, transcriptional profiles, inherited traits, hormonal landscape, brain structure and function, as well as treatment responses to antidepressants ^2,5–7^. Nevertheless the specific cellular and molecular origins of sex bias remain unclear ^8^, hindering the development of sex-specific biomarkers for early diagnosis and sex-tailored therapies.

Oligodendroglial lineage cells, including OPCs and mature oligodendrocytes (OLs), play multifaceted roles in the central nervous system (CNS), including forming myelin sheaths to support axonal function and metabolism, as well as conducting myelin independent functions such as synaptic communication with neurons and regulation of immune responses ^9,10^. Among these important functions, myelin deficits have been reported in patients with MDD, as reflected by reduced white matter integrity ^11,12^. In addition, single nucleus transcriptomic data have identified aberrant OPCs as a potential link to MDD ^13–15^. It is known that neuronal activity promotes the myelination to meet the connectomics demands of neuronal networks ^16^; it remains however unclear how OPCs development and myelination are coordinated with neuronal states in MDD, particularly under sex-biased conditions. While, a limited number of studies have begun to investigate sex-specific regulatory mechanisms centered on the oligodendroglial lineage in depression ^17,18^, the intercellular communication networks that involve oligodendroglia and contribute to the increased vulnerability to depression observed in women, have not been fully explored.

To address this intercellular communication network, current approaches primarily rely on clustering methods for analyzing differentially expressed genes (DEGs). However, these methods overlook subtle contributor genes that fall below detection thresholds, and often miss contributions arising from interactions among multiple factors, rather than the isolated effect of individual components. Interactions between various cell types in the brain are shaped by distinct cell-cell communications ^19^, with complex networks mediating environmental adaptation and maintaining homeostasis. Although some studies have incorporated network analyses, such as protein-protein interactions (PPIs), but the lack of cell-type specificity limits the ability to observe the dynamics of population-level signaling. Moreover, decontextualized gene or protein lists fail to capture the directional causality of signaling, particularly sender-receiver asymmetry, which is critical for understanding intercellular communication ^20^. As a result, sex-specific intercellular communication in MDD remains a black box.

In the present study, we established a framework that integrates cutting-edge computational analyses of human patient data with histological and biochemical validation to identify sex-specific mechanisms in MDD. By deciphering system-level, sex-biased dysregulation within interactome networks, we identified within oligodendroglial lineage ^10^ a subpopulation of OPCs, as key preferential signal receivers. The NRXN synaptic assembly pathway was the most perturbed, with female-specific exacerbation. In OPC subpopulation, the γ-aminobutyric acid type A receptor subunit γ3 (*GABRG3*) gene emerged as an MDD-specific hub gene, which is also involved in the NRXN cascade, mechanistically linking the molecular deficits to depression through neuron-OPC synaptic communication. Using conditional knockout mice, we validated findings in patients, by demonstrating that conditional deletion of *Gabrg3* in OPCs recapitulates MDD-like phenotypes, including impaired neuron-OPC synaptic communication, abnormal myelination, and depression-like behaviors. Furthermore, this approach enabled us to determine potential underlying mechanisms. Our work demonstrates a sex-specific cellular and molecular pathobiology of MDD, bridges clinical pathology and preclinical discovery, and offers a translatable framework for future sex-tailored interventions.

## Results

### 1. Sex-specific dysregulation of cell-cell interactome networks in MDD

We conducted a sex-stratified comparative dissection of intercellular communication networks in MDD by systematically analyzing curated human single-nucleus RNA sequencing (snRNA-seq) datasets from the dorsolateral prefrontal cortex (dlPFC) of both sexes ^14,21^, focusing on identifying sex-biased pathological alterations, and subsequently confirmed these human-derived findings using traditional mouse models, thereby offering cross-species validation and highlighting previously unrecognized, sex-specific intercellular signaling in MDD pathology **(Supple. Fig. 1)**.

To quantify the network-wide impairment, we figured the percentage of alteration in global interaction strength which was calculated as the communication probability by CellChat ^22^ for females and males separately. Our results indicated a predominant female-biased global attenuation, with signal reductions of 23.04% (*p* = 0.031) in females compared to 4.1% (*p* = 0.02) in males **(Fig. 1A)**, which connects with the epidemiological observations that women exhibit higher incidence rate of MDD compared to men ^2^. We next integrated these communication networks with the largest sex-stratified MDD genome-wide association studies (GWAS) ^6^ and quantified the enrichment of the derived genes across all cell types. We found that the oligodendroglial lineage cells exhibited a significantly stronger cell-type-specific association with these sex-biased MDD risk genes, exceeding that observed in neurons and other neuroglial cells **(Fig. 1B)**, which may suggest divergent roles of distinct cells in the sexually dimorphic pathophysiology of MDD. In line with this, we plotted the interaction strength across individual cell clusters within four cohorts separately **(Fig. 1C)**, and revealed a marked heterogeneity in communication patterns across different cell types in MDD, with distinct alterations observed between female and male MDD patients (*p* < 0.0001, **Fig. 1C)**.

**Figure 1.**
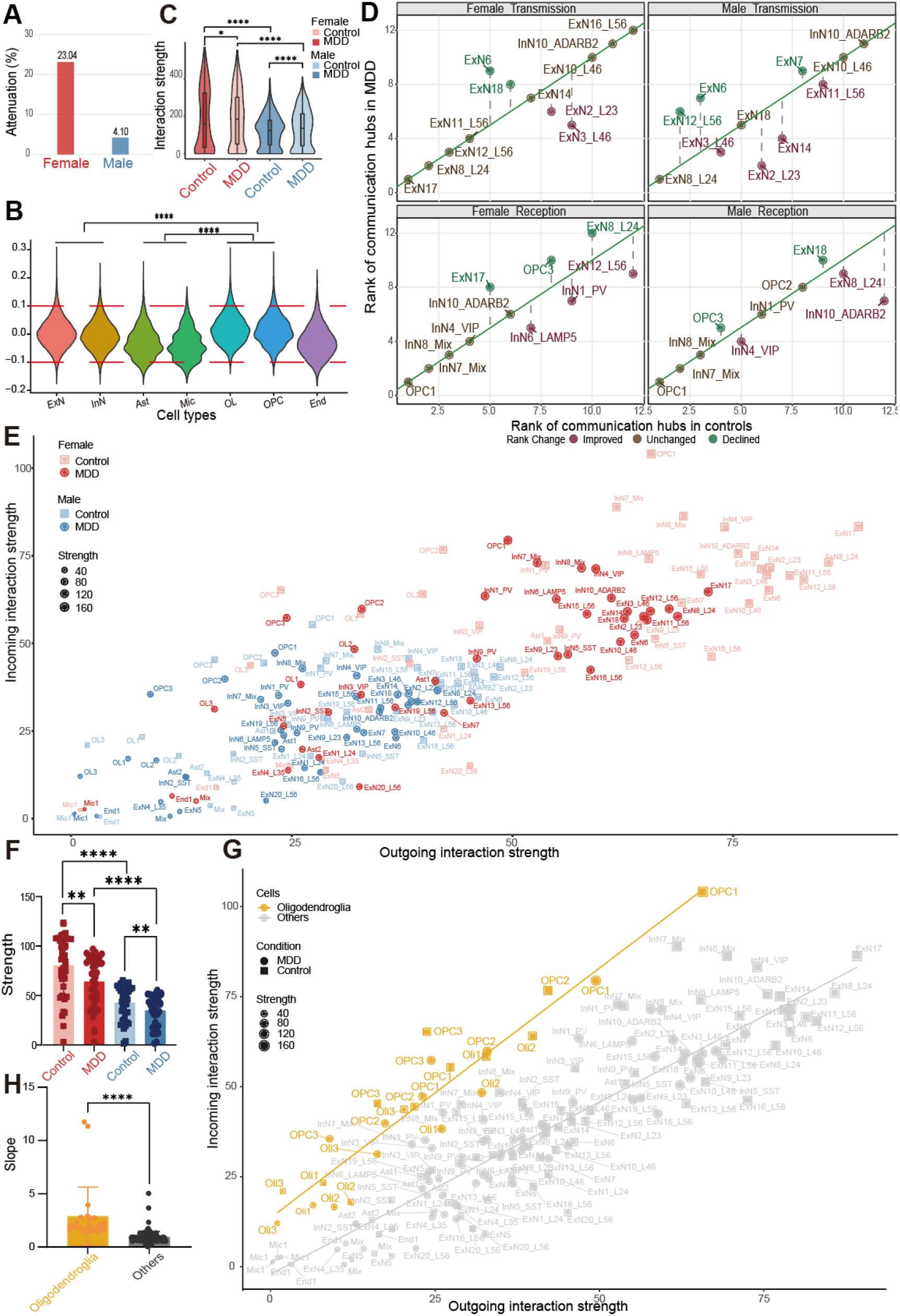
Mapping sex-stratified cell-cell interactome dysregulation in MDD. (A) Percentage of alteration in global interaction strength for female and male MDD cohorts compared to controls. (B) Cell-type-specific enrichment of sex-stratified MDD GWAS risk genes. (C) Violin plots showing the distribution of interaction strengths across individual cell clusters within four cohorts (Female Control, Female MDD, Male Control and Male MDD). (D) Rank concordance of top communication hub clusters between MDD patients and controls within each sex. (E) Scatter plot showing the transmission-reception propensity of distinct cell clusters across four cohorts. The x-axis and y-axis represent the interaction strength for signal transmission (outgoing) and reception (incoming). (F) Bar plot quantification of global interaction strength across four cohorts. (G) Regression mapping demonstrating that oligodendroglial lineage cells form a high-slope (reception versus transmission) boundary population compared to all other cells. (H) Bar plot displaying the deviation in transmission-reception propensities for oligodendroglial lineage cells compared to all others. Statistical significance for (A) was determined via permutation analysis. Differences in (B), (C), (F) and (H) were evaluated using the two-sided Kolmogorov-Smirnov test in GraphPad Prism. \**p* < 0.05, \*\**p* < 0.01, \*\*\**p* < 0.001, \*\*\*\**p* < 0.0001.

To investigate how sex bias in MDD arises from sex-specific topological vulnerabilities within distinct cell clusters, we compared heatmap signatures of intercellular interaction strengths **(Supple. Fig. 2A, 2B)**. Although both sexes exhibited widespread reductions in signaling strength (blue spots) across cell types, female MDD networks were characterized by highly concentrated and intense focal interactions, whereas male networks displayed diffuse and disorganized connectivity landscape **(Supple. Fig. 2A, 2B)**. Analyses of the top communication hubs revealed a clear sexually dimorphic pattern of network disruption **(Fig. 1D)**. Specifically, female MDD networks showed highly preserved signaling transmission but severely disrupted reception. In contrast, males exhibited impaired transmission alongside coherent reception. Of note, OPC subsets (notably OPC1 subset) consistently retained their status as dominant receivers in both sexes, highlighting the vulnerability of OPCs in depression **(Fig. 1D)**.

In addition, we implemented a bidirectionality matrix mapping the sender-receiver propensities across cellular populations **(Fig. 1E)**. Female cell communities occupied peripheral coordinates with a greater Euclidean distance from origin, reflecting enhanced dual-directional engagement in both efferent and afferent signals (points with red and pink hues). Male networks exhibited limited interactome flexibility, with the majority of populations confined to the bottom-left quadrant along the central diagonal (points with navy and blue hues). Furthermore, the global attenuation of interaction strength within dysfunctional networks associated with MDD was characterized by a noticeable retraction of cells in MDD groups across sexes (illustrated by darker hues in **Fig. 1E**; and shorter bars in **Fig. 1F**). Remarkably, oligodendroglial lineage cells deviated from the diagonal, forming a high-slope boundary population (slope = 1.387 vs. other cells’ slope = 0.956, *p* < 0.0001), emphasizing their unidirectional receptivity bias **(Fig. 1G, 1H; Supple. Fig. 2C)**.

Overall, our results indicate that female-preserved bidirectional plasticity contrasts with male-constrained interactome rigidity, while suggesting oligodendroglial lineage as a potential pathological integrator in MDD progression.

### 2. Aberrant synaptic connectivity in female MDD patients

To comprehend the communication signatures linked to sex-specific pathophysiological network in MDD, we assessed the divergence in interactions between MDD and controls for each cell cluster by calculating Euclidean, horizontal (x-axis), and vertical (y-axis) distances **(Supple. Fig. 3A)**. Notably, female networks exhibited a greater overall signaling capacity and a more pronounced attenuation in communication intensity, characterized by steep and progressively stabilized declines in interaction gradients, in contrast to the gradual moderation observed in males **(Supple. Fig. 3A)**. In line with that, we categorized the top 10 most divergent cell clusters (populated by neurons and oligodendroglial lineages cells) in three dimensions for each sex **(Fig. 2A)**, subsequently quantifying the proportional representations of these two lineages within their respective broad populations. Once again, the results highlighted the functional prominence of neurons and oligodendroglial lineages cells in the network dynamics associated with MDD and a distinct bias of oligodendroglial lineage cells in signal reception **(Fig. 2A, 2B; Supple. Fig. 3B)**.

**Figure 2.**
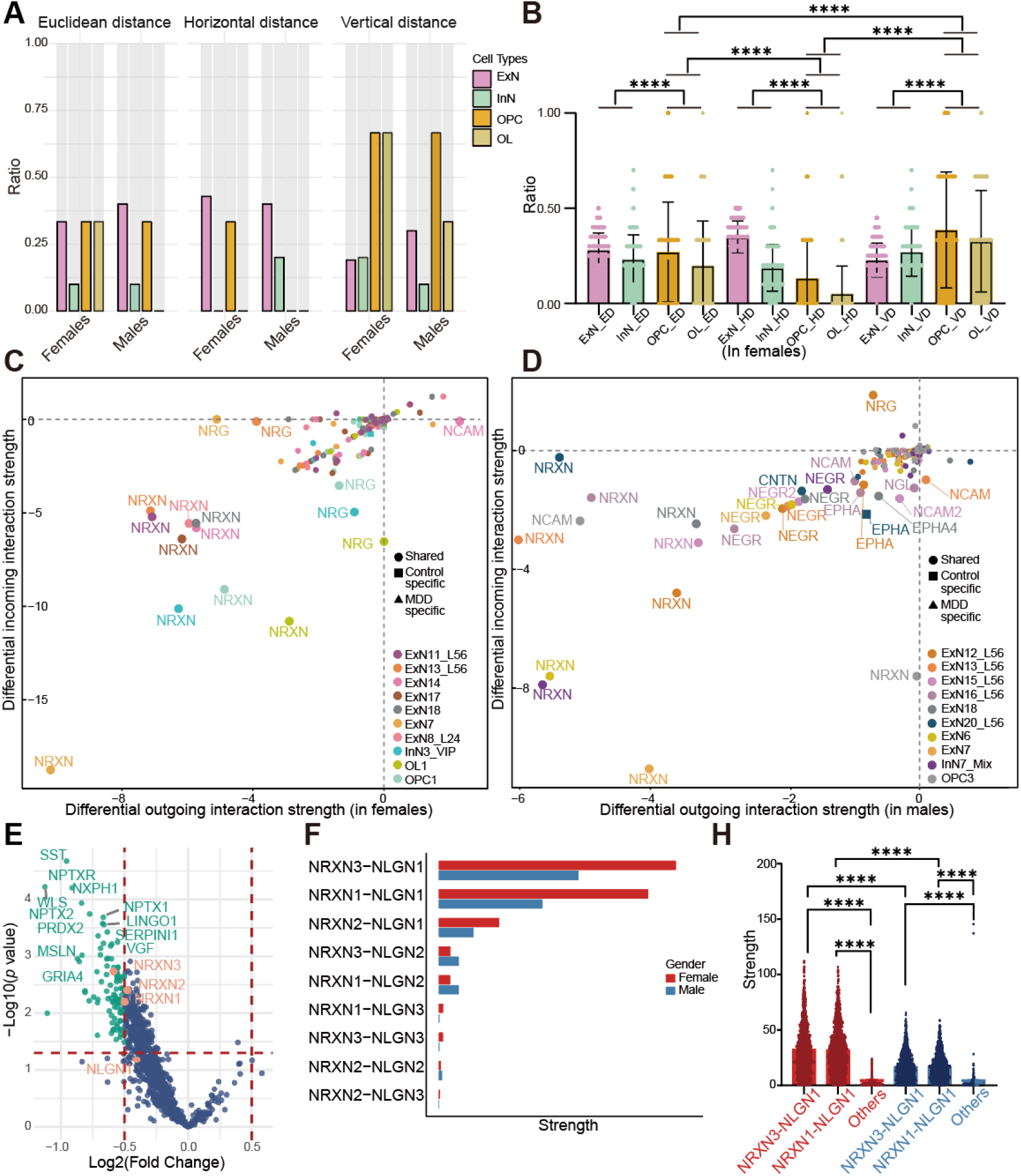
Synaptic dysconnectivity and NRXN pathway perturbation in MDD. (A) Categorization of the top 10 most divergent cell clusters between MDD patients and controls in three dimensions across both sexes. (B) Proportional representations of the top 10 most divergent cell clusters in females, highlighting the distinct functional bias of oligodendroglial lineage cells in signal reception. (C-D) Plots visualizing differential outgoing and incoming signaling changes of the top 10 most divergent cell clusters from control to MDD in (C) females and (D) males. (E) Clinical evaluation displaying the levels of NRXN3 and NRXN1 proteins detected in the cerebrospinal fluid of depression patients. (F) Comparative analysis showing the signaling changes of dysregulated ligand-receptor pairs from control to MDD within the NRXN pathway in both sexes. (G) Plot demonstrating NRXN3-NLGN1 and NRXN1-NLGN1 as the most dysregulated interaction pairs in both sexes. Data distributions for (B), (F) and (H) were generated via permutation analysis, and subsequent statistical significance between groups was evaluated using the two-sided Kolmogorov-Smirnov test in GraphPad Prism. \**p* < 0.05, \*\**p* < 0.01, \*\*\**p* < 0.001, \*\*\*\**p* < 0.0001.

In addition, pathway complexity analysis revealed a higher multiplicity of dysregulation in females, with female MDD networks engaging 12 perturbed pathways (*p* = 0.025) compared 4 in males (*p* = 0.753; **Supple. Fig. 3C**), suggesting a greater level of attenuation and complexity in the pathophysiology of MDD among females. Notably, the NRXN (Neurexin) pathway was the most affected pathway in MDD, and its disturbance was significantly more severe in females **(Fig.2C, 2D; Supple. Fig. 3C).** In addition, the specific signaling changes of the top 10 most divergent cell clusters within the NRXN network all localized to the extremes in quadrant III, corresponding to a broad reduction in both signal transmission and reception **(Fig.2C, 2D)**. The dysfunctional NRXN pathway was further confirmed by the relatively lower levels of NRXN3 and NRXN1 proteins in the cerebrospinal fluid of depression patients ^23^ **(Fig. 2E)**. Additionally, comparative analysis of ligand-receptor dyads in MDD patients revealed NRXN3-NLGN1 (neuroligin1) and NRXN1-NLGN1 as the most impaired interaction pairs linked to NRXN pathway suppression in both sexes **(Fig. 2F)**. The more pronounced signal alterations observed in these pairs in females highlights higher vulnerability of synaptic connectivity in women with MDD **(Fig. 2F, 2G)**.

Together, our findings identify disrupted oligodendroglial signal reception as a core feature of female MDD, further implicating NRXN pathway-mediated synaptic dysconnectivity as the principal pathological alteration.

### 3. Sexually dimorphic roles of OPCs in synaptic malfunction in MDD

To decode the specific cellular and molecular alterations of oligodendroglial lineage cells in MDD pathogenesis, we applied NeuronChat, a neural-specific cell-cell communication inference tool, which incorporates neurotransmitter signaling, to map intercellular signaling networks ^24^. We found that OPCs exhibited the most prominent baseline activity in signal reception (indicated by bright red spots in the frames) across all cohorts **(Supple. Fig. 4)**. Comparative heatmap of interaction strengths in MDD patients versus healthy controls revealed a markedly pronounced reduction in communication probabilities in females (globally darker blue hues), compared to the milder deficits observed in males **(Fig. 3A, 3B)**. Specifically, OPCs displayed a substantial attenuation in receiving signals from neurons in females **(Fig. 3A)**, in contrast to that of a comparatively modest malfunction of OPCs in males **(Fig. 3B)**. Meanwhile, mature OLs exhibited distinctly different and subtler alteration profile, lacking profound reception deficit **(Fig. 3C)**. This stark functional asymmetry within the lineage firmly demonstrates OPC-centric communication breakdown as the primary mechanism underlying the sex-specific disease manifestations.

**Figure 3.**
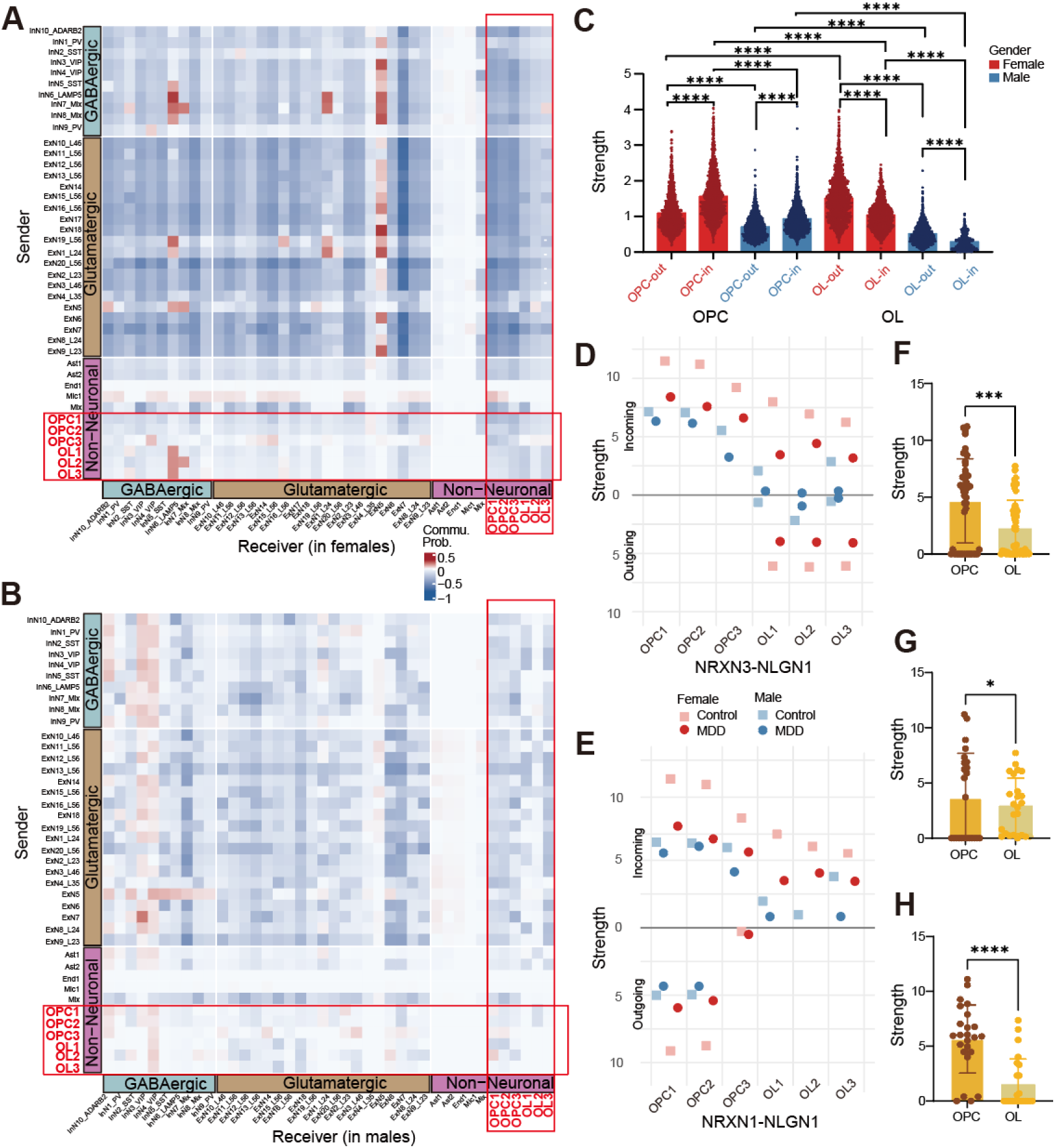
Sex-divergent mapping of oligodendroglial communication interactions. (A-B) Heatmaps displaying the differential neural-specific interaction strengths among cell clusters between controls and MDD in (A) females and (B) males. Red (or blue) spots represent increased (or decreased) signaling in MDD cohort compared to controls. (C) Plot demonstrating the differential alteration profiles within the oligodendroglial lineage cells. (D-E) Transmission-reception mapping of interaction capacities of the oligodendroglia for the specific (D) NRXN3-NLGN1 and (E) NRXN1-NLGN1 ligand-receptor pairs. A greater distance from the x-axis along the y-axis indicates higher signaling intensity. (F-H) Statistical quantifications comparing the interaction stability between OPCs and OLs. This includes (F) the overall variance across both NRXN ligand-receptor pairs, and the specific engagement capacities for (G) NRXN3-NLGN1 and (H) NRXN1-NLGN1 pairs. Data distribution for (C) was generated via permutation analysis, followed by group comparisons using the two-sided Kolmogorov-Smirnov test. Statistical significance for (F), (G) and (H) were directly evaluated utilizing the two-sided Kolmogorov-Smirnov test in GraphPad Prism. \**p* < 0.05, \*\**p* < 0.01, \*\*\**p* < 0.001, \*\*\*\**p* < 0.0001.

Building on the suppression of the NRXN pathway in MDD (as shown in Fig. 2), we further investigated how oligodendroglia contribute to neuron-OPC synaptic dysconnection, specifically their sex-divergent signaling patterns underlying molecular malfunction. By integrating the interactions from the single NRXN3-NLGN1 and NRXN1-NLGN1 ligand-receptor pairs within the NRXN pathway **(Supple. Fig. 5)**, we demonstrated that OPCs and OLs exhibited competitive capacities for signaling reception rather than transmission (data points primarily positioned toward the positive y-axis; **Fig. 3D, 3E)**. OPCs displayed a broader amplitude variance along the x-axis, reflecting significantly greater signaling dynamism than OLs (*p* = 0.0002; **Fig. 3D, 3E, 3F**). Moreover, beyond signal intensity, OPCs and OLs engaged distinct repertoires of the prioritized NRXN ligand-receptor pairs (NRXN3-NLGN1 pair: *p* = 0.013; **Fig. 3G**; NRXN1-NLGN1 pair: *p* < 0.0001; **Fig. 3H**).

Collectively, our findings identify OPCs as pivotal mediators of the sex-dimorphic network dysfunction in MDD, linking cellular communication biases to molecular pathway disruptions thus offering a unified framework for understanding the heterogeneity of this disease.

### 4. Oligodendroglial Ident3 emerges as a central pathogenetic hub for neuron-OPC synaptic dysregulation in female MDD

To uncover mechanistic links between OPCs and dysregulation of neuron-OPC synapses, we performed unsupervised clustering of the whole oligodendroglial population (OPC1, 2, 3, and OL1, 2, 3), and identified 3 distinct cellular subsets (Ident 1, Ident 2 and Ident 3) visualized in the UMAP plot **(Fig. 4A)**. Notably, Ident 1 was predominantly composed of OPCs, Ident 2 was enriched with OLs, while Ident 3 exhibited a mixed composition of both OPCs and OLs as indicated by a hybrid coloration of cells **(Fig. 4A)**. Using CellChat analysis, we quantified the communication probabilities of NRXN-NLGN pairs across these Idents. Neuron-to-OPC interactions predominantly occurred in OPC-containing clusters Ident 1/3 and through NRXN3-NLGN1/NRXN1-NLGN1 pairs (darker red spot indicates the higher probability) **(Supple. Fig. 6A)**; whereas OPC-to-neuron interactions likely occurred via the NRXN1-NLGN1 pair (only a few spots in the frame) **(Supple. Fig. 6B)**, once again, highlighting NRXN pathway-mediated synaptic dysconnectivity as the key pathological alteration in Idents.

**Figure 4.**
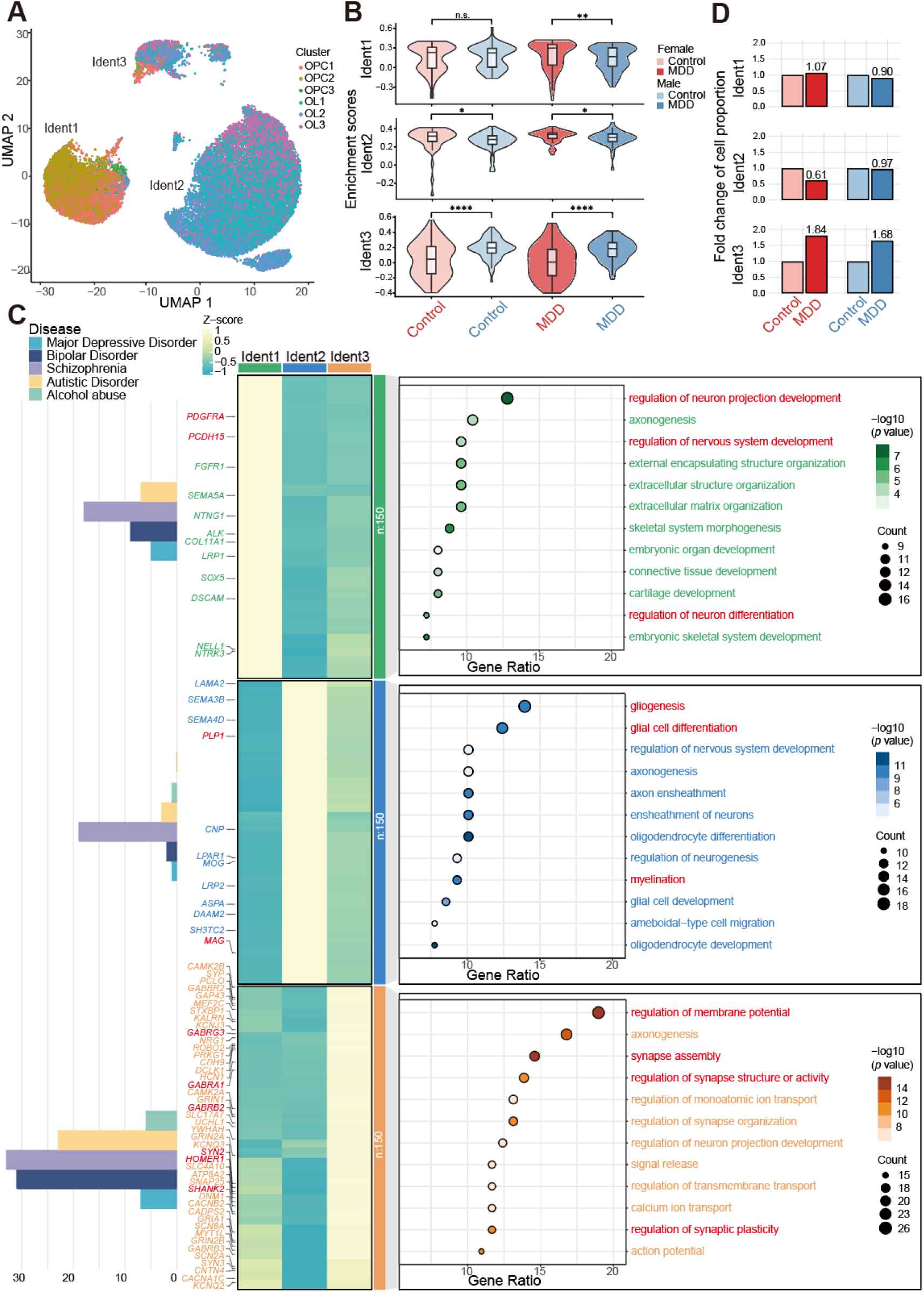
Cellular and molecular characterization of pathogenic oligodendroglial subsets in MDD. (A) UMAP plot displaying the three subsets of oligodendroglial lineage cells. (B) Violin plots evaluating the expression profiles of ident-specific 150 markers across four cohorts within each subset. (C) Comprehensive functional characterization of the oligodendroglial subsets. Left: bar plots illustrating the quantitative disease relevance scores from DisGeNET ^25^; middle: heatmaps depicting the expression signatures of distinct marker genes across the subsets; right: dot plots of the GO enrichment analysis characterizing the functional divergence of the subsets. (D) Bar graphs displaying the normalized fold change for the cell proportions within three subsets. Controls are normalized to 1.0 and MDD cohorts are represented as the relative proportional alterations. Statistical significance for (B) was evaluated using the two-sided Kolmogorov-Smirnov test in GraphPad Prism. n.s. not significant, \**p* < 0.05, \*\**p* < 0.01, \*\*\**p* < 0.001, \*\*\*\**p* < 0.0001.

We further calculated the enrichment of the Ident-specific top 150 signature genes across the four cohorts for each subset, revealing that Ident 3 exhibited the most profound sex-specific transcriptomic divergence among the three subsets **(Fig. 4B)**. Subsequently, we applied DisGeNET (Disease Gene NETwork ^25^), a curated repository of genes and variants linked to human diseases, to conduct a quantitative analysis of disease relevance. Bar plots **(Fig. 4C; left panel)** illustrated that Ident3 had the strongest disease relevance, with the highest number of enriched genes (100 in total) among neuropsychiatric disorders, including MDD, bipolar disorder, schizophrenia, autistic spectrum disorders, and alcohol abuse. Heatmaps **(Fig. 4C; middle panel)** highlighted key markers, such as *PDGFRA* (Platelet Derived Growth Factor Receptor α), *PCDH15* (Protocadherin 15) in Ident1, as well as *PLP1* (Proteolipid Protein 1) and *MAG* (Myelin Associated Glycoprotein) in Ident2, suggesting the identities of Ident1 as a progenitor-like state, and Ident2 as a mature OL-like state. The gene ontology (GO) enrichment analysis **(Fig. 4C; right panel)** further indicated that Ident1 was functionally enriched in developmental processes including the regulation of neural projection development, nervous system development and neuron differentiation, whereas the Ident2 is centered on gliogenesis, neuroglia differentiation and myelination. Furthermore, Ident3 was characterized by its markers relevant to mental disorders **(Fig. 4C; middle panel)**, such as *SHANK2* (SH3 and Multiple Ankyrin Repeat Domains 2), and synaptic regulation, including *HOMER1* (Homer Scaffold Protein 1), *SYN2* (Synapsin II), *GABRA1* (γ-Aminobutyric Acid Type A Receptor Subunit α1), *GABRB2* (γ-Aminobutyric Acid Type A Receptor Subunit β2), and *GABRG3* (γ-Aminobutyric Acid Type A Receptor Subunit γ3). Once again, these results suggest a strong involvement of Ident3 in membrane potential regulation, synapse assembly, regulation of synapse structure and activity and synaptic plasticity **(Fig. 4C; right panel)**, directly linked to MDD pathophysiology^26,27^.

We next sought to quantify disease-driven cellular shifts within oligodendroglial subsets by calculating fold changes in cell proportions in MDD patients relative to sex-matched healthy controls **(Fig. 4D)**. In Ident 2, we observed a consistent reduction in proportion of OLs in both male and female MDD patients, aligning with MDD-associated demyelination ^28^. Notably, female patients exhibited a more pronounced reduction than males, suggesting greater loss of mature OLs and a more severe myelin deficit in females **(Fig. 4D)**. Conversely, Ident 3 proportions were markedly increased in MDD patients of both sexes, further supporting Ident 3 as the subset most strongly associated with the disease state. Moreover, this pathological expansion was relatively higher in females, pointing to an aberrant cellular response predominantly to female MDD patients **(Fig. 4D)**. This concurrent loss of Ident 2 and expansion of Ident 3 in females reveal a female-biased pathological divergence in oligodendroglial dynamics.

Collectively, our results reveal oligodendroglial Ident3 are the central hub cells for female MDD pathogenic activity, suggesting that synaptic dysregulations in MDD may originate from these cells.

### 5. GABRG3-bearing committed OPCs drive sex-divergent neuron-OPC synaptic dysregulation in MDD

The subclustering analysis of entire Ident3 revealed three transcriptionally distinct subclusters, which were ordered along the oligodendroglial differentiation pseudotime trajectory: OPCs, committed OPCs (cOPCs), and mature OLs **(Fig. 5A)**. Cellular identities were confirmed through visualization of the canonical cell-specific markers, namely *PDGFRA* and *CSPG4* (Chondroitin Sulfate Proteoglycan 4 or NG2) for OPCs, *ITPR2* (Inositol 1,4,5-Trisphosphate Receptor Type 2) and *FYN* (FYN Proto-Oncogene, Src Family Tyrosine Kinase) for cOPCs, and *PLP1* (Proteolipid Protein 1) and *MOG* (Myelin Oligodendrocyte Glycoprotein) for OLs **(Supple. Fig. 6C)**.

**Figure 5.**
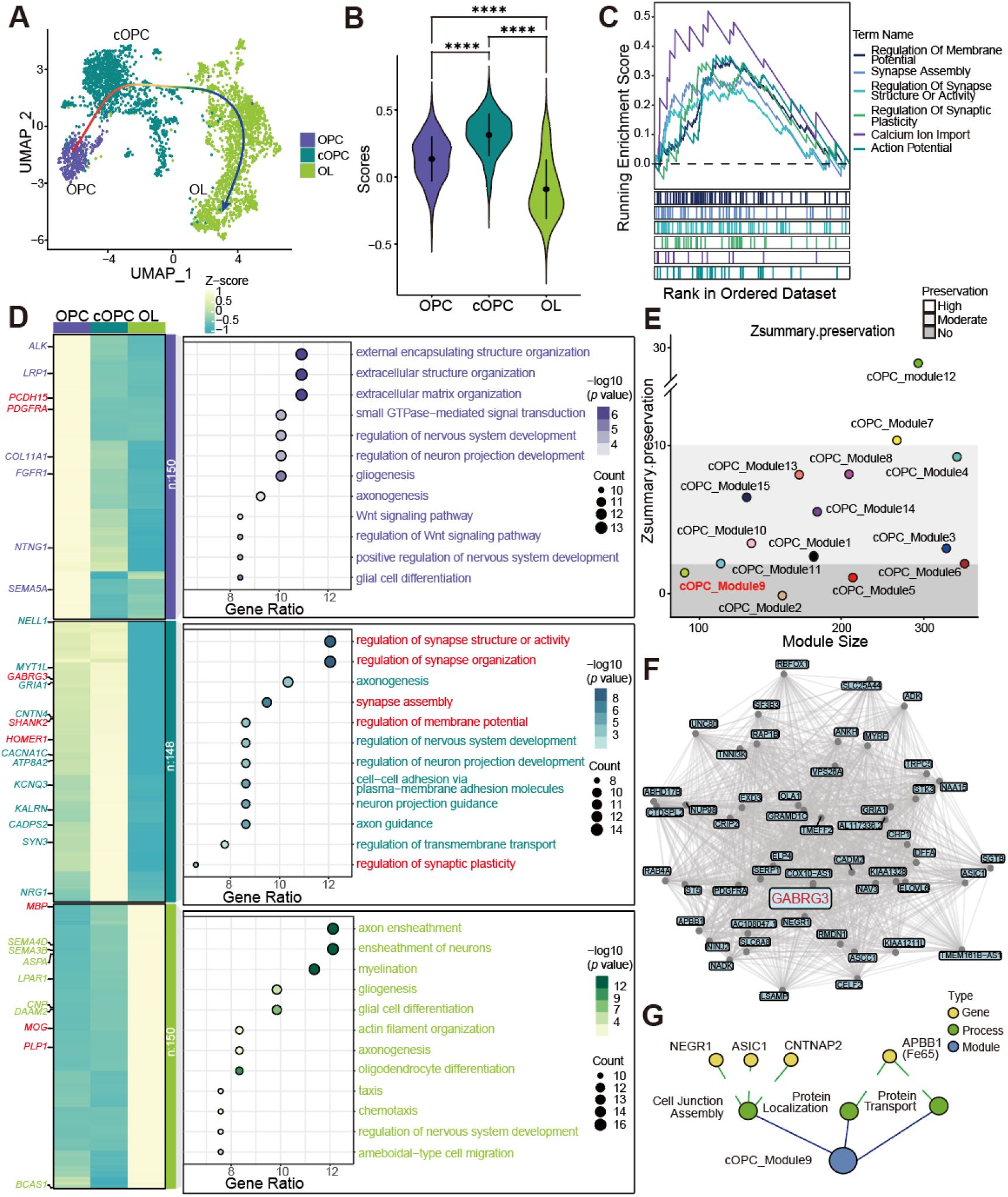
Transcriptional subclustering and co-expression network characterization of *GABRG3*+ committed OPCs. (A) UMAP plot displaying the subclustering of Ident3 population along a pseudotime trajectory. (B) Violin plots evaluating the enrichment scores of the top 150 transcriptional signatures of Ident3 across identified subclusters. (C) GSEA plots evaluating the activity of gene sets associated with synaptic pathways within the cOPCs. (D) Distinct marker genes and GO enrichment analysis characterizing the functional specificity across the identified subclusters. (E) Scatter plot displaying the preservation (Z-summary score) of co-expression modules identified from healthy control cOPCs in MDD. Preservation levels are demonstrated by three different shades: no (dark grey, Z < 2), moderate (2 < Z < 10), and high (Z >10). (F) Network plot validating *GABRG3* as the hub gene within the MDD-specific subnetwork of cOPC_Module9. (G) Schematic plot highlighting the established synaptic modulators and cell adhesion molecules within the *GABRG3*-centered subnetwork of cOPC_Module9. Statistical significance for (B) was evaluated using the two-sided Kolmogorov-Smirnov test in GraphPad Prism. \**p* < 0.05, \*\**p* < 0.01, \*\*\**p* < 0.001, \*\*\*\**p* < 0.0001.

To determine which subclusters primarily contribute to sex-divergent pathological responses, we assessed the enrichment of the MDD-related top Ident3-specific 150 transcriptional signatures across the three identified subclusters. We found that cOPCs displayed significantly higher enrichment of this specific gene set compared to OPCs and OLs **(Fig. 5B)**. Additionally, we conducted GSEA (Gene Set Enrichment Analysis) to evaluate the activity of the gene sets in cOPCs associated with GO terms of neuronal excitability and synaptic signaling. This analysis again confirmed that genes responsible for processes regulating neuronal membrane potential, synapse assembly, and synaptic plasticity, were collectively and significantly enriched in cOPCs **(Fig. 5C)**, reinforcing the role of cOPCs as central hub cells in modulating MDD pathophysiology. GO enrichment analysis further validated that the key pathways such as the regulation of synapse structure or activity and synapse organization, regulation of membrane potential, synapse assembly, and synaptic plasticity, which significantly enriched in Ident3 were convergently active in cOPCs, but were largely absent in conventional OPCs and OLs **(Fig. 5D)**.

In order to further identify the potential target gene(s) that underpin the molecular mechanism of cOPCs in MDD pathophysiology, we employed hdWGCNA (high-dimensional Weighted Gene Co-expression Network Analysis ^29^), a computational framework designed for analyzing co-expression networks based on single-cell transcriptomics data. By projecting 15 identified modules from healthy controls into the MDD cohort, we evaluated the preservation and quality statistics of these modules via Z-summary scores **(Fig. 5E; Supple. Fig. 6D)**. Specifically, modules in the preservation plot were stratified into three distinct shades (white, light gray, and dark gray), forming a gradient that reflects a functional spectrum. Modules in the white zone are highly conserved across conditions, while those in the dark gray zone show no preservation in MDD, thus representing the highest specificity to MDD.

As shown in **Fig. 5E**, cOPC_Module9, cOPC_Module2, and cOPC_Module5 fell into the dark gray zone, indicating their highest specificity to MDD. Notably, although all three modules may contribute to MDD pathophysiology, cOPC_Module9 was distinguished by the inclusion of *GABRG3* among its hub genes, which is among the most prioritized transcriptional signatures in Ident3 **(Fig. 5F; Supple. Fig. 6E)**, whereas cOPC_Module2 and Module5 lacked association with Ident3. To further analyze the functionalities of *GABRG3*-centered subnetwork of cOPC_Module9, we conducted GO analysis on the enriched genes in this module. Our results revealed that the cOPC_Module9 was functionally characterized by the biological processes including regulation of cell junction assembly, protein localization and protein transport **(Supple. Fig. 6F)**. Critically, the key effector genes within this module (*NEGR1*, Neuronal Growth Regulator 1, *ASIC1*, Acid Sensing Ion Channel Subunit 1, *CNTNAP2*, Contactin Associated Protein 2, and *APBB1*, Amyloid-β Precursor Protein Binding Family B Member 1) are all established players in synaptic modulation and cell adhesion **(Fig. 5G)**, indicating that *GABRG3* emerges not merely as a receptor gene, but as the topological and functional hub of this phenotype-aligned subnetwork in MDD.

Given prior evidence ^30–33^ implicating *GABRG3* in the NRXN3-NLGN1-gephyrin-GABRG3 synaptic assembly signaling cascade in neurons, our results suggest that this *GABRG3*-mediated synaptic assembly pathway in cOPCs contributes to a molecular dysfunctional network underlying sex-divergent neuron-OPC synaptic dysconnectivity in MDD **(Supple. Fig. 6G)**. Thus, *GABRG3* emerges as a candidate for functional validation.

### 6. Expression patterns of GABRG3-mediated synaptic assembly signaling in mouse cOPCs

To validate the role of *GABRG3*-mediated synaptic assembly signaling in MDD, we firstly investigated the expression patterns of *Gabrg3*, along with *Nrxn3*, *Nlgn1* and *Gephyrin* in various cell types in the CNS. Although *Gabrg3* mRNA is highly expressed by multiple cell types in the CNS, including both OPCs and neurons, as shown by the UMAP plot from mouse whole-brain transcriptomic cell type atlas ^34^, *Gabrg3* is the top listed GABA receptor subunit gene expressed in OPCs **(Fig. 6A, 6B)**. Meanwhile, mRNA of *Nrxn3*, *Nlgn1* and *Gephyrin* was also present in both OPCs and neurons **(Fig. 6C)**.

**Figure 6.**
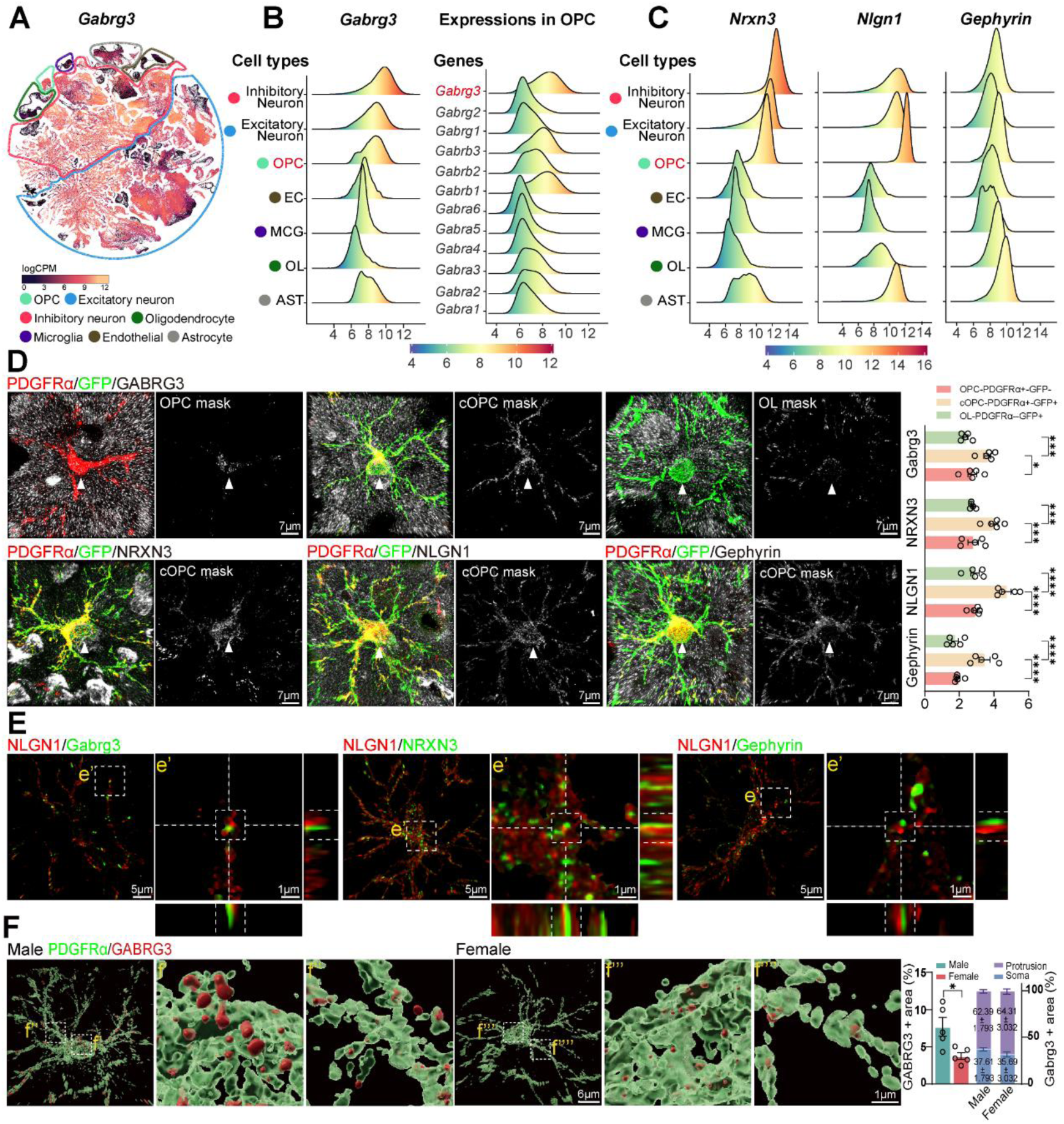
Expression patterns of synaptic elements in oligodendroglia. (A) UMAP plot showing *Gabrg3* gene expression across different cell types of whole mouse brain from the Allen Brain Cell Whole Mouse Brain Atlas. (B) Ridgeline plot showing *Gabrg3* gene expression across different cell types of mice brain (left), and different GABR subtype genes in OPCs of mice brain (right). Gene expression data was acquired from the Allen Brain Cell Whole Mouse Brain Atlas. (C) Ridgeline plots showing the *Nrxn3, Nlgn1* and *Gephyrin* gene expressions across different cell types of whole mouse brain from the Allen Brain Cell Whole Mouse Brain Atlas. (D) Representative images of PDGFRα, Tau-GFP and synaptic elements staining in oligodendroglial lineage cells during their developmental process by super-resolution fluorescence microscopy. The number of synaptic elements per unit of oligodendroglial lineage + area. Scale bar = 7 µm, n=5 mice. (E) Representative images of synaptic elements in cOPCs by super-resolution fluorescence microscopy. Orthogonal view of the square area indicates the co-expressions of these synaptic elements in cOPCs. Scale bar = 1 µm and 5 µm, n=5 mice. (F) Surface rendering images of GABRG3 expression in mice. Quantifications indicate the GABRG3+ area per unit PDGFRα+ area, as well as its protein localization to both cellular processes and the soma. Scale bars = 1 µm and 6 µm; n = 5 mice. Data presented as mean ± SEM; \**p* < 0.05, \*\*\**p* < 0.001, \*\*\*\**p* < 0.0001. Also see Supplementary Figure 7.

We further examined the protein levels of GABRG3, NRXN3, NLGN1 and gephyrin in oligodendroglial lineage cells during their development by super-resolution fluorescence microscopy. *NG2*^CreERT^;*Tau-mGFP* mouse strain was employed, in which mGFP gene is under control of the Tau promoter upon tamoxifen induction ^35^. Besides, SOX10 antibody was used to label whole oligodendroglial lineage, PDGFRα antibody was used to label OPCs, and IP_3_R-II antibody was used to label cOPCs ^36^. Generally, combinations of these markers indicate that SOX10+/PDGFRα+/IP_3_R-II-/mGFP-cells were OPC, SOX10+/PDGFRα+/IP_3_R-II+/mGFP+ cells were cOPCs, and SOX10+/PDGFRα-/IP_3_R-II-/mGFP+ cells were newly generated mature OLs **(Supple. Fig. 7A)**. The GABRG3, NRXN3, NLGN1 and gephyrin synaptic elements were observed on the surface or closely associated with OPC’s cell body and main processes, as shown by the 3D-reconstructions **(Fig. 6D; Supple. Fig. 7B)**. These synaptic elements were significantly upregulated in PDGFRα+/mGFP+ cOPCs when compared to other OPCs and mature OLs **(Fig. 6D; Supple. Fig. 7B)**. In addition, as shown by super-resolution fluorescence microscopy and 3D-reconstructures, after positive signal masked by PDGFRα signal using Imaris, the co-expressions of these synaptic elements in cOPCs were verified **(Fig. 6E; Supple. Fig. 7C)**. We further examined sex differences in the protein levels of these synaptic elements. It appeared that GABRG3 protein exhibited significant sex-divergent expression **(Fig. 6F; Supple. Fig. 8A)**. In contrast, other proteins showed no sex-related differences, nor did their subcellular localization differ between sexes **(Fig. 6F; Supple. Fig. 8A)**. These findings suggest that sex-specific synaptic dysconnection in MDD is likely to arise from disease-associated modulation.

### 7. OPC-specific *Gabrg3* knockout recapitulates MDD-like pathophysiology across behavioral, cellular, and molecular levels in a transgenic mouse model

To mimic the female-biased neuron-OPC synaptic dysconnection in MDD, we newly generated a *Gabrg3^fl^*^/fl^ mouse strain and crossed it with the OPC-specific *PDGFRα*^CreER^ mouse to achieve conditional *Gabrg3* knockout mice **(Fig. 7A)**. Since *Gabrg3* encodes a GABA receptor subunit, the primary inhibitory neurotransmitter in the mammalian brain, the accumulated vGAT signal in inhibitory synapses were decreased by 33.77±7.02%, but the accumulated vGlut1 signal in excitatory synapses were not altered **(Fig. 7B)**.

**Figure 7.**
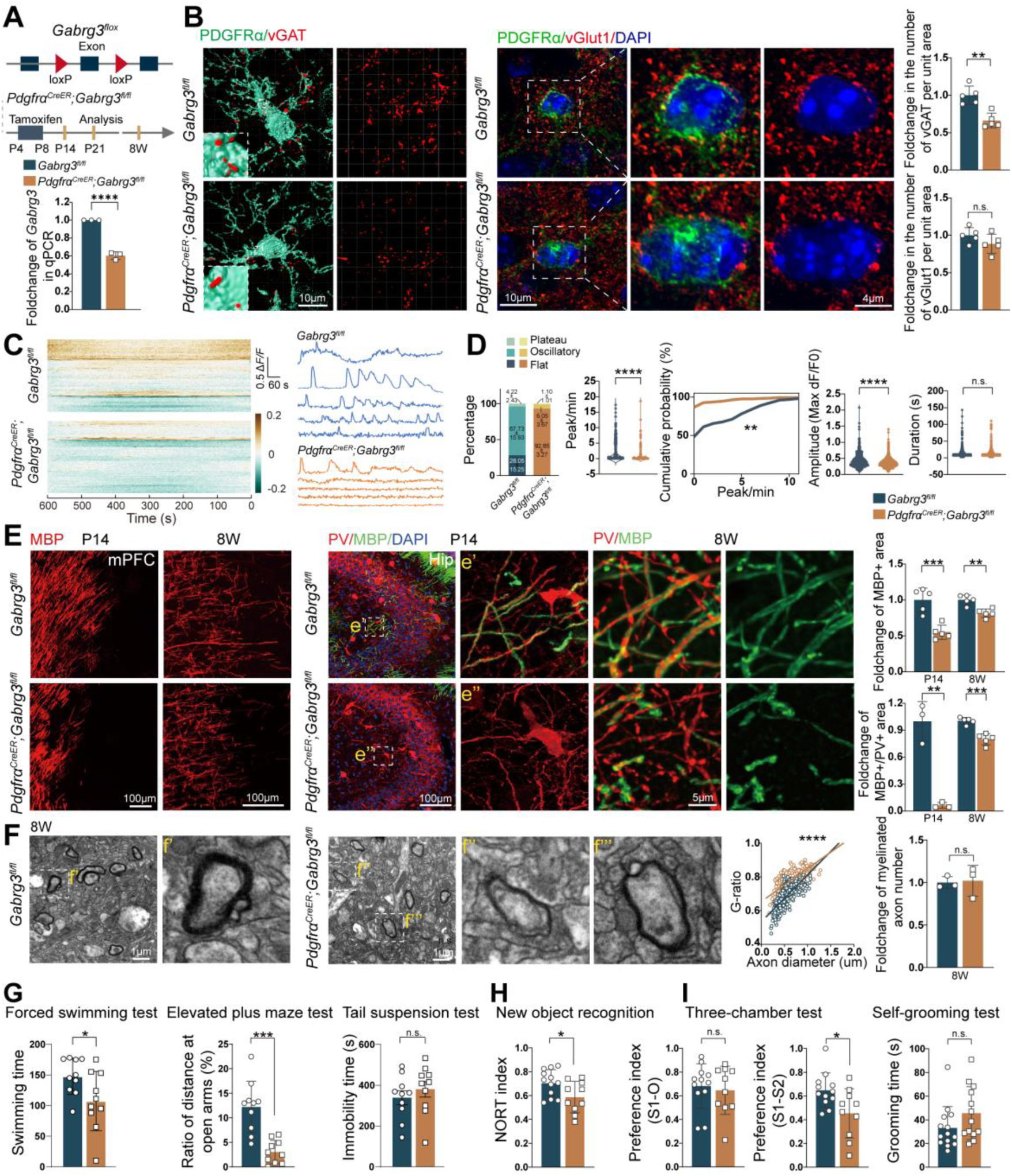
OPC-specific *Gabrg3* knockout induces MDD-like pathophysiology at behavioral, cellular, and molecular levels. (A) Establishment of *PDGFRα*^CreER^; *Gabrg3*^fl/fl^ conditional knockout mouse strain and the experimental diagram. mRNA expression of *Gabrg3* gene in isolated OPCs from *Gabrg3* knockout and control mice by immunopanning. n = 3 independent experiments. (B) Surface rendering images of PDGFRα/vGAT staining in the mPFC of *PDGFRα*^CreER^; *Gabrg3*^fl/fl^ and control mice with at P14 using Imaris software. Immunofluorescence staining of PDGFRa/vGLUT1 in the mPFC from *PDGFRα*^CreER^; *Gabrg3*^fl/fl^ and control mice at P14. Quantifications indicate the number of vGAT, vGLUT1 per unit of PDGFRα+ area. Scale bar = 10 µm, n = 5 mice. (C-D) Representative recording of cytoplasmic Ca^2+^ dynamics OPCs in brain slices; real-time intracellular Ca^2+^ in OPCs with different conditions are shown respectively (C); the quantification (D) of OPC Ca^2+^ images in *PDGFRα*^CreER^; *Gabrg3*^fl/fl^ and control mice. (E) Representative images of MBP staining in the mPFC of *PDGFRα*^CreER^; *Gabrg3*^fl/fl^ and control mice at P14 and 8W. Scale bar = 100 µm, n = 5 mice. Representative images of MBP and PV staining in the hippocampal CA3 region of *PDGFRα*^CreER^; *Gabrg3*^fl/fl^ and control mice at P14, and SMI32, PV and MBP staining in the hippocampal CA3 region at 8W. Scale bar = 100 µm, 5 µm. Upper panel quantification indicates MBP+ areas in the mPFC at different developmental stages. Lower panel quantification indicates MBP+ area in the PV+ region at P14, n = 3 mice, and MBP+ area in the SMI32+ / PV+ region at 8W, n = 5 mice. (F) Electron microscopy image of the mPFC and quantification of myelinated axon number and g-ratio at 8W. Scale bar = 1 µm, n = 3 mice. (G-I) Behavioral test results of (G) forced swimming test, elevated plus maze test and tail suspension test, (H) new object recognition, (I) three-chamber test and self-grooming test in *PDGFRα*^CreER^; *Gabrg3*^fl/fl^ and control mice. Data presented as mean ± SEM; \**p* < 0.05, \*\**p* < 0.001, \*\*\**p* < 0.001, \*\*\*\**p* < 0.0001. Also see Supplementary Figure 8.

To examine the neuron-OPC synaptic communication in *Gabrg3* deletion mice, we measured the intracellular Ca^2+^ dynamics in PDGFRα+ OPCs in acutely isolated brain slices. Consistent with our previous study, there were three types of intracellular Ca^2+^ dynamic patterns: (i) “flat” Ca^2+^ signaling; (ii) spontaneous “oscillatory”- Ca^2+^ signaling with peaks, and (iii) spontaneous high “plateau” transients. In *Gabrg3* deletion mice, most OPCs showed “flat” Ca^2+^ dynamics, while a few OPCs exhibited spontaneous oscillatory-like Ca^2+^ changes and with peak/plateau Ca^2+^ signals **(Fig. 7C)**, reflecting an abnormal response of OPCs lacking *Gabrg3* to neuronal activity. We also determined the characteristics of Ca^2+^ dynamics in OPCs. The number of peaks and the amplitude were significantly decreased in *Gabrg3* deficient OPCs **(Fig. 7D)**, whereas the duration of Ca^2+^ responses was similar to that in wildtype OPCs **(Fig. 7D)**. These findings indicate that loss of *Gabrg3* in OPCs impairs their responses to neuronal activity.

Consistent with the view that neuron-OPC synaptic connection regulates OPC differentiation and myelination ^36^, we observed myelin defects in *Gabrg3* knockout mice, as reflected by reduced MBP-positive areas across serval depression-related brain regions, specifically the mPFC and hippocampus, but not in the striatum, corpus callosum, cerebellum, lateral habenula, or habenula **(Fig. 7E; Supple. Fig. 8B)**. These myelin deficits also persist in development though diminishing with time as shown for mPFC at P14 and 8 weeks of age **(Fig. 7E)**. In particular, the reduction of myelination rate of axons of parvalbumin (PV)-positive inhibitory interneurons in the hippocampus was gradually changed from a decrease of 94.13±12.96% at P14, to a decrease of only 20.43±3.22% at 8 weeks of age **(Fig. 7E)**. Myelin thickness, as shown by electron microscopy, significantly decreased in mPFC at 8 weeks of age, though the myelinated axon numbers didn’t show significant difference at that time point **(Fig. 7F).**

We next determined whether *Gabrg3* deletion in OPCs could lead to the onset of MDD-like symptomology. We assessed behavioral outcomes related to MDD. *Gabrg3* knockout mice showed a marked increase in immobility time in the forced swimming test, as well as a strong tendency toward increased immobility in the tail suspension test **(Fig. 7G)**. In the elevated plus maze, *Gabrg3* deficient mice exhibited a significant reduction in distance traveled in the open arms **(Fig. 7G)**, thus indicating that loss of *Gabrg3* leads to depressive-like behavior. Moreover, *Gabrg3* knockout mice displayed a range of additional behavioral alterations. In the novel object recognition test, they showed marked memory alteration **(Fig. 7H)**. In the three-chamber test, the *Gabrg3* knockout mice exhibited social novelty deficits **(Fig. 7I)**; however, no changes were observed in the self-grooming test **(Fig. 7I)**. These behavioral changes suggest that *Gabrg3* knockout mice are susceptible to depressive-like mood changes, memory impairments, and social novelty deficits.

Thus, we validate that the targeted alteration of a specific postsynaptic gene *Gabrg3* can lead to cOPC-neuron synaptic communication deficit, and subsequent myelin abnormalities, as well as depressive-like behavioral outcomes in mice.

### 8. Transcriptomic profiling reveals shared and distinct differentially expressed genes and pathways in *Gabrg3* knockout OPCs

To further elucidate the possible molecular links between depression-related symptoms and the observed demyelination phenotype in both the mPFC and hippocampus, we performed RNA-seq on the purified OPCs from these two regions of *Gabrg3^fl^*^/fl^ and *PDGFRα*^CreER^; *Gabrg3*^fl/fl^ mice **(Fig. 8A)**. Differential expression analysis revealed profound transcriptomic changes in both regions **(Fig. 8B)**. Notably, both regions exhibited a significant upregulation of estrogen receptor 1 (*Esr1)* triggered by the loss of GABAergic synaptic input. Specifically, in the mPFC OPCs, the upregulated genes were predominantly enriched for extracellular matrix (ECM) components and structural regulators (e.g., *Col1a2*, *Col3a1*, *Ptgs2*, *Epha4*, *Mc4r*, *Draxin*, *Postn* and *Gper1*), and collagen proteins have been reported to impair OPC differentiation and myelination ^37^. At the same time, downregulated transcripts included those directly associated with early neurodevelopment and synaptic connectivity (e.g., *Gabra1*, *Nrn1*, *Gad2*, *Dlx1/2* and *Arx*) **(Fig. 8B; left panel)**. Hippocampal OPCs displayed a striking downregulation of crucial glial homeostatic sensors (*Spp1*, *Gria1*, *Itgam*, *Mertk*, *Cd36*, *Lgals3*, *Pirb, Trem2*, *Tyrobp*, *Aqp4*) **(Fig. 8B; right panel).** Hippocampal OPCs also exhibited an upregulation of ECM deposition and established markers relevant to mental disorders (*Col1a1*, *Col8a1*, *Dcn*, *Nlrp3*, *Penk*, *Serpinf1*, *Serpine1*).

**Figure 8.**
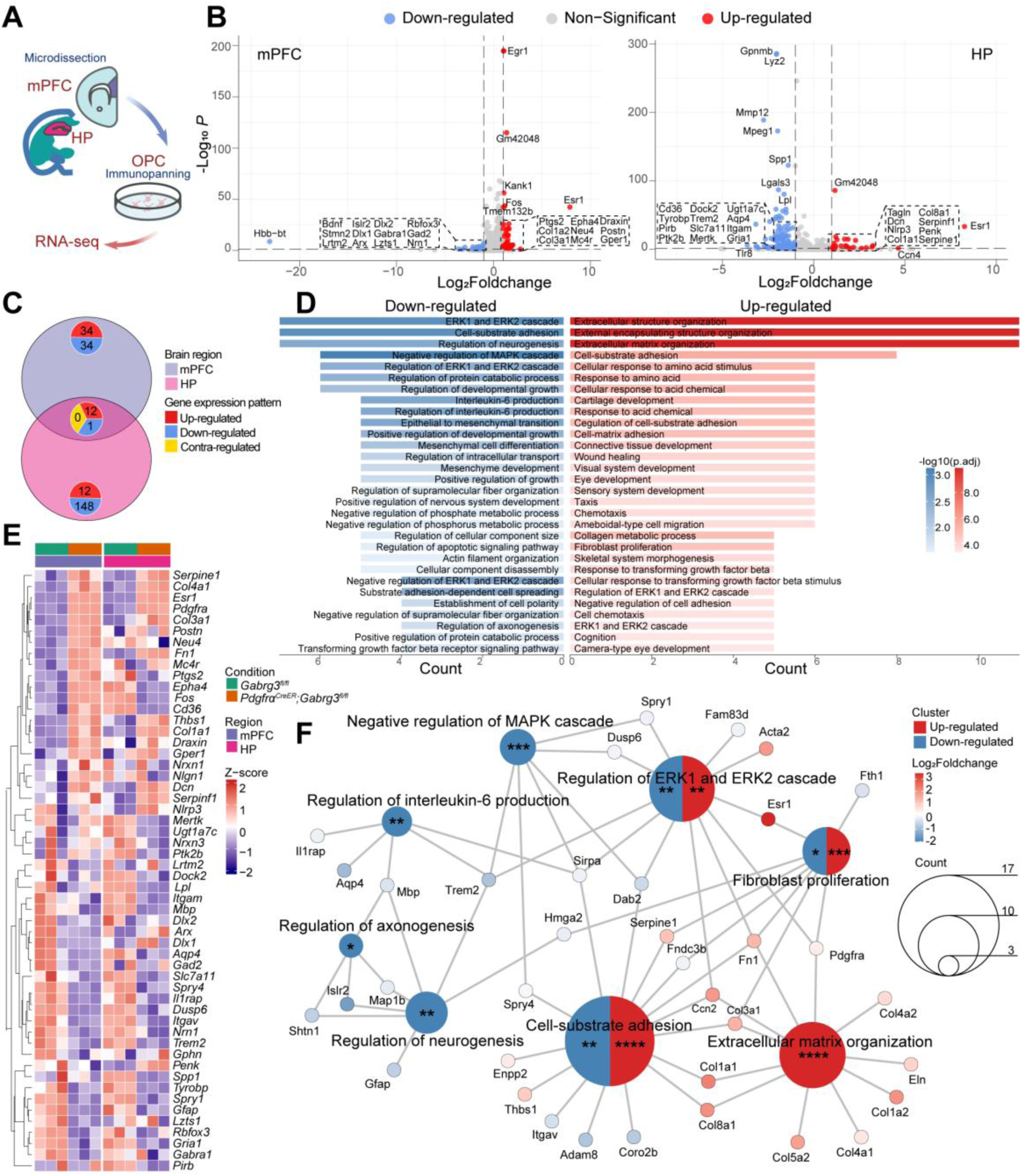
Transcriptomic profiling of Gabrg3-deficient OPCs reveals conserved disruption of maturation signaling and extracellular matrix remodeling. (A) Diagram of the experimental workflow: purifying OPCs by immunopanning from the mPFC and hippocampus of *Gabrg3*^fl/fl^ and *PDGFRα*^CreER^; *Gabrg3*^fl/fl^ mice for RNA-seq. n = 3 independent experiments. (B) Volcano plots depicting differentially expressed genes (DEGs) in purified OPCs from the mPFC (left) and hippocampus (right) of *PDGFRα*^CreER^; *Gabrg3*^fl/fl^ versus control mice. Significantly up-regulated (log_2_Foldchange >1, red) and down-regulated (log_2_Foldchange <1, blue) genes with adjusted *p*-value < 0.05 are highlighted. (C) Venn diagram illustrating the overlap of consistently up- (red) and down-regulated (blue) genes between the mPFC and hippocampal OPCs. (D) GO (BP) enrichment analysis of the convergently up- (red) and down-regulated (blue) gene sets (with adjusted *p*-value < 0.05). Top 30 significantly enriched pathways are shown, ranked by the number of associated genes (count). (E) Heatmap showing the normalized expression (Z-score) for the curated panel of genes across all biological replicates. (F) Integrated pathway-gene network model summarizing the proposed mechanism. The significance of enriched pathways was denoted as: \**p* < 0.05, \*\**p* < 0.01, \*\*\**p* < 0.001, \*\*\*\**p* < 0.0001.

Venn diagram analysis was utilized to compare the differentially expressed genes between the two regions **(Fig. 8C)**; GO enrichment analysis indicated that the commonly upregulated genes in both regions were predominantly enriched in processes driving structural microenvironment remodeling, including ECM organization, cell-substrate adhesion and fibroblast proliferation **(Fig. 8D)**. Conversely, the commonly downregulated genes were exclusively enriched in pathways critical for OPC differentiation and myelination, specifically the regulation of MAPK and the ERK1/ERK2 cascade, neurogenesis and axonogenesis **(Fig. 8D)**.

We further selected a representative set of core genes to construct a mechanistic model of the *Gabrg3* deletion-driven processes **(Fig. 8E)**. The chosen genes included key drivers such as the sex hormone receptor (*Esr1*), essential OPC lineage receptors (*Pdgfra*), major ECM depositors (*Col* family), critical regulators of the intracellular MAPK pathway (*Dusp6*, *Spry1*/*4*), as well as markers of glial dysfunction and myelination failure (*Trem2*, *Aqp4*, *Mbp*) **(Fig. 8E)**. Visualization of their normalized expressions across all biological replicates revealed a highly consistent regulatory pattern in both brain regions **(Fig. 8E)**. Finally, the pathway-gene network analysis illustrated the crosstalk among these functional modules **(Fig. 8F)**.

The network provides a conceptual overview of our proposed overarching mechanism: loss of *Gabrg3*-mediated GABAergic input from neurons triggers an aberrant state in cOPCs, and ultimately leading to impaired myelination.

## Discussion

In the present study, we established a framework that integrates computational analyses of human patient data with animal models of MDD; furthermore, we uncovered pronounced sex-specific vulnerabilities that shape the disease trajectory. Specifically, we identified a sexually divergent synaptic assembly signaling involving NRXN3, NLGN1, and most prominently GABRG3, which orchestrates neuron-OPC crosstalk and drives the pathogenesis of MDD in females.

Many neurological diseases exhibit pronounced sex differences. Women are more susceptible to anxiety, depression and Alzheimer’s disease, while men are more prone to autism and Parkinson’s disease ^4,38^. The MDD in particular displays notable sex-biased prevalence ^4^. Early bulk RNA-sequencing studies of brain tissues reveal marked alterations in GABA-related genes within the prefrontal cortex (PFC) of female but not male, mice ^39^, supporting our notion that the GABA gene network is specifically altered in women with depression ^39^. With the advent of single-cell RNA-sequencing, sex differences in cellular and molecular signatures across specific brain cell types became evident ^14,21,39^, implicating glutamatergic/GABAergic transmission, glial malfunction, and blood-brain barrier integrity ^14^. Since then, multiple brain cell types have been linked to MDD pathogenesis. Recently, several computational tools and algorithms that decode cell-cell interactions at single-cell resolution were developed ^19^. In the present study, we integrated multiple computational approaches, including intercellular signaling inference, trajectory reconstruction, gene co-expression networking, and disease-association mapping, to decode cell type-specific interactome, dissect sex-specific intercellular signaling networks, and track the functional adaptation of OPCs during disease progression. After comprehensively quantifying and evaluating these networks from MDD patient single-cell transcriptomic data, we validated our findings in traditional animal experimental methods, a step often lacking in bioinformatic studies.

Traditional MDD research has primarily focused on neuronal dysfunction, viewing glial cells as passive supporters ^40^. However, postmortem studies multiple glial abnormalities in MDD ^41^, including reduced glial numbers in the PFC and hippocampus, altered astrocyte morphology, microglial reactivity, and oligodendrocyte changes ^7,41–43^. Single-nucleus transcriptomics of treatment-resistant depression further implicates glial-specific genes, including a hub gene, *EAAT2* (encoding Excitatory amino acid transporter 2, critical for glutamate homeostasis in the CNS ^44^); the deficiencies in the latter contributes to pathogenesis of several disorders of cognition ^45^. OPCs, as the precursors of oligodendroglia in the CNS ^46^, exhibit functions beyond myelination ^10,36,47^. Transcriptomic data from MDD patients show reduced expression of OPC differentiation and glutamatergic transmission regulating genes ^48^. Deep-layer excitatory neurons and OPCs accounted for 47% of transcriptional changes observed in the dorsolateral PFC of males with MDD ^14^. In animal studies from our laboratory and others, OPC depletion in mice induces depression-like behaviors in response to stress ^49,50^. Yet, no study ever elucidated sex-specific mechanisms of OPC-driven MDD pathogenesis. Here, our computational analyses reveal disrupted neuron-OPC synaptic communication and identify committed OPCs as a central cellular subpopulation in MDD responsible for sex bias. Our findings are further supported by a recent study ^51^ that mapped sex differences in gene expression in the human cerebral cortex at single-cell resolution, identifying over 3,000 genes with differential expression levels between male and female. Among these, oligodendroglia showed the strongest sex-differential effects, which may explain why certain neurological diseases linked to myelin damage or white matter abnormalities (including MDD), display pronounced sex biases in incidence or pathological presentation.

As a critical pathophysiological feature, myelin deficits, including reduced myelin content, OL density, OPC differentiation, and myelin gene expression, were observed in emotion-related brain regions such as the medial prefrontal cortex, hippocampus, amygdala, and corpus callosum of patients with MDD ^28^. Female patients, particularly, exhibit greater myelin deficits in corpus callosum and hippocampus ^28^. In line with this, preclinical data indicate that juvenile stress reduces OL density and myelin only in adult females ^52^, and that restorative treatments effective in males have lesser effect in females ^53^. Some clinical findings, however, are mixed: using R1 (inverse of T1 signal, sensitive to myelin), female adolescents with MDD showed higher R1 than healthy controls ^54^, whereas adults (19-58 years) with MDD exhibited lower global R1 ^54^. These observations provide a clinical interpretation for our findings: female-biased loss of oligodendroglial Ident2 reflects chronic mature OL loss as a consequence of MDD; more importantly, the expansion of disease-associated Ident3 reiterates an exacerbated dysconnectivity of neuron-OPC synapses in females, which leads to pathological OPC differentiation arrest, blocks the transition from cOPC to mature OL, and failed myelin formation/turnover in female MDD.

Given the critical role of neuron-OPC synapses in shaping OPC development and myelination, as supported by our studies and those of others ^36,47,55^, the changes in their component, assembly signaling and downstream pathway in MDD remain unclear. This molecular complexity hindered the development of diagnostic biomarkers and therapeutic targets. Although previous hypotheses implicated biogenic amines, neurotransmitter receptors, neurotrophic factors, cytokines, and hormones ^26^, none of these can fully explain MDD’s multifaceted nature. Transcriptome studies reveal sex-divergent synapse-related gene expressions in MDD patients (downregulated in males, upregulated in females) ^7,56^. In the present study, pathway deconvolution revealed disrupted NRXN signaling in cOPCs from female MDD patients. NRXNs and NLGNs are synaptic adhesion proteins linked to neuropsychiatric disorders ^33^, with different NRXN-NLGN pairs serving distinct synaptic roles ^57^. NLGN1 and NLGN2 are brain-specific and bind gephyrin ^57^. Although glial NRXN/NLGN expression has been reported, showing that females exhibit increased *NRXN1* expression associated with AD and MS in oligodendroglia ^58^, and *NLGN1* is upregulated in oligodendroglia for remyelination in female MS patients ^59^, our study is the first to show NLGN1 colocalization with gephyrin and GABRG3 in cOPCs, distinct from neurons ^31,60^, albeit the underlying mechanism remains unknown. Regarding GABA signaling, GABA acts on ionotropic GABA_A_ and metabotropic GABA_B_ receptors with well-defined neuronal roles ^61^. GABA receptor expression in oligodendroglia is confirmed ^61^, but their roles in OPC differentiation are debated. For instance, GABA_A_ receptors, including GABRG3, decrease with maturation and inhibit differentiation/myelination, whereas GABA_B_ receptors remain stable and promote these processes ^62,63^. We speculate that the distinct functional outcomes of GABAergic signaling may vary by brain region, developmental stage, and specific GABA receptor subtype. In *Gabrg3* knockout OPCs, *Esr1* was significantly upregulated. Its higher expression and link to increased female neuron numbers ^64^ suggest this upregulation may reflect a sex-biased compensatory mechanism when OPCs lose neuronal connection. Beyond *Esr1*, the expression profiles of other genes in *Gabrg3* knockout OPCs exhibited regional heterogeneity, which may help connect our preclinical findings to the broad spectrum of symptoms observed in patients. These diverse symptoms could be driven by commonly upregulated genes that are involved in microenvironment remodeling, along with commonly downregulated genes associated with OPC differentiation and myelination^65^. This process could potentially occur through the MAPK and ERK1/ERK2 cascades, which are known to play key regulatory roles in myelination ^66^, GABA_A_ receptor function ^67^, and sex-dependent antidepressant treatment ^68,69^.

We acknowledge the limitations of this study. The role of GABRG3 in cOPC-mediated synaptic signaling alterations in MDD remains unclear, and the identified synaptic genes in cOPCs are not exclusive to female MDD, making it difficult to specifically target pathological cOPCs or cOPC-driven neural activity without affecting neurons or other physiological functions. Although evidence implicates myelin integrity alterations in MDD, the precise mechanisms underlying impaired myelin deficits during MDD pathogenesis remain elusive, hindering myelin-oriented therapy development. Looking ahead, as generative artificial intelligence continues to evolve ^70^, purpose-driven digital research on mental disorders may enable more effective exploration of underlying mechanisms and treatment strategies, ultimately transforming the field of mental health research.

In summary, our work reveals a sex-specific cellular and molecular pathobiology in female MDD, bridges preclinical discovery and clinical pathology by identifying a neuron-OPC synaptic communication hub linked to depressive symptoms, and may offer an integrative computational-animal framework for future mechanistic and therapeutic research.

## Materials and methods

### Acquisition of the human single-nucleus RNA-sequencing dataset

Single-nucleus RNA sequencing (snRNA-seq) datasets of the dorsolateral prefrontal cortex (dlPFC) from human MDD patients and healthy controls were acquired from NCBI GEO database under accession numbers GSE144136 and GSE213982 ^14,21^, composed of 37 MDD patients (20 females, 17 males) and 34 matched healthy controls (18 females, 16 males).

### Sex-stratified network comparison and topological analysis

To infer intercellular communication networks, we utilized the R packages CellChat ^22^ and NeuronChat ^24^. The overall network attenuations were quantitatively assessed to quantify the network-wide impairment in both sexes. A sex-stratified MDD genome-wide association studies (GWAS) ^6^ was integrated to detect the cell-type-specific association with these sex-biased MDD risk genes. Network centrality was performed and revealed a sexually dimorphic pattern of network disruption. The aggregated outgoing and incoming interaction strengths of each cell cluster across four cohorts were projected onto 2D scatter plots to map the sender-receiver propensities, and three spatial distance metrics were computed to reflect their distinct extent of pathological deviation. Comparative analysis of the disturbance of pathways and corresponding ligand-receptor pairs uncover the aberrant communication signatures in MDD.

### Pathological and functional interpretation of oligodendroglia

The oligodendroglial lineage cells were computationally isolated and subjected to high-resolution subclustering. Three primary functional subsets (Ident 1, Ident 2, and Ident 3) were identified, characterized by distinct transcriptomic profiles and canonical marker expression. With the list of ident-specific top 150 transcriptomic signatures, Ident3 showed the strongest disease relevance reported in DisGeNET ^25^. Cell proportion fold change analysis was conducted to quantify disease-driven cellular shifts within oligodendroglial subsets. Furthermore, the cells of Ident 3 were extracted for secondary refined subclustering, which revealed three subclusters along the inferred pseudotime trajectory. In addition, high-dimensional weighted gene co-expression network analysis (hdWGCNA ^29^) was performed to identify co-expression modules and evaluate their preservation across conditions.

### RNA-sequencing and analysis

RNA-seq was performed on the *Gabrg3* knockout OPCs from both the mPFC and hippocampus. Differential expressed genes and enriched pathways were compared in order to elucidate the potential molecular links between depression-related symptoms and the observed demyelination phenotype.

### Gene expression analysis from online mouse scRNA-seq platform

The Mouse Whole-Brain Transcriptomic Cell Type Atlas was accessed through the Allen Institute for Brain Science portal (https://portal.brain-map.org) ^34^. The platform offers us the data to plot the expression of *Gabrg3*, *Nrxn3*, *Nlgn1*, *Gephyrin* and various Gabr subtypes across different cell types.

### Mice

All animal studies were performed under the guidelines of laboratory animal welfare and ethics committee of the Third Military Medical University (AMUWEC20223048). All mice were housed in a temperature- and humidity-controlled environments with free access to standard chow and water and on a 12 h/12 h light/dark cycle. Gabrg3-flox mice were generated by Gempharmatech Co.,Ltd. (China). The mice were then crossed with *PDGFRα*^CreER^ mice to generate *PDGFRα*^CreER^; *Gabrg*3^fl/fl^ conditional KO mice. NG2^CreERT^; Tau-mGFP mice were used to label the newly formed OLs.

### Behavior tests

Mice were handled daily for five days before the test. The apparatus was wiped with 80% alcohol between each trial. All tests were conducted b**et**ween 9 am and 6 pm. The experimenters were blinded for the grouping. VisuTrack Animal Behavior Analysis Software (Shanghai XinRuan) was used for data collection and analysis. Depression-related behavior was evaluated using the tail suspension test and the forced swimming test. Anxiety-like behavior was measured using the elevated plus maze and the open field test. Sociability and preference for social novelty were evaluated using the three-chamber test. The self-grooming test assessed repetitive behaviors in mice. Novel object recognition test evaluated short-term memory function.

### Statistical analysis

GraphPad Prism 9.0 (GraphPad Software, San Diego, CA, USA) was used to determine statistical significance. Data are presented as mean ± standard error of the mean (SEM). Comparisons between two experimental groups were made using unpaired two-tailed t-tests. P-values < 0.05 were considered statistically significant, with significance levels indicated as: \**p* < 0.05, \*\**p* < 0.01, \*\*\**p* < 0.001, \*\*\*\**p* < 0.0001. Data distribution was assumed to be normal, although this was not formally tested. Sample sizes were not predetermined by statistical methods but were comparable to those reported in previous studies. Each experiment was repeated at least three times.

## Acknowledgements

This work was supported by grants from National Natural Science Foundation of China (W2511095 to J.N. and A.V.; 32271034 to J.N.). Chongqing Natural Science Fund for Distinguished Young Scholars (CSTB 2023NSCQ-JQX0030 to J.N.).

## Data availability

Human snRNA-seq datasets are from GSE144136 and GSE213982 ^14,21^. Mouse OPC RNA-seq data in this manuscript has been deposited in the NCBI GEO database under the accession number GSE334265. Any additional information required to reanalyze the data reported in this paper is available from the lead contact upon request.

## Code availability

All scripts used to analyze data are provided in a Github repository (https://github.com/leeyinong/MDD_data_analysis) and a Zenodo repository (https://zenodo.org/records/20535058).

## Author contributions

J.N. conceived the study. J.N. A.V. and C.Z. designed the experiments. C.Z. and Y.L. performed bioinformatic analyses. X.W., Z.W., Y.S., S.L., S.W., X.C., and L.W. performed the animal experiments and analyzed the data. A.V. and Y.W. contributed to discussion. J.N., A.V. and C.Z. wrote the manuscript. A.V. made the editing. J.N. is the lead contact.

## References

1. Fava, M., and Kendler, K.S. (2000). Major Depressive Disorder. Neuron 28, 335–341. 10.1016/S0896-6273(00)00112-4.

2. Eid, R.S., Gobinath, A.R., and Galea, L.A.M. (2019). Sex differences in depression: Insights from clinical and preclinical studies. Progress in Neurobiology 176, 86–102. 10.1016/j.pneurobio.2019.01.006.

3. Seney, M.L., Glausier, J., and Sibille, E. (2022). Large-Scale Transcriptomics Studies Provide Insight Into Sex Differences in Depression. Biological Psychiatry 91, 14–24. 10.1016/j.biopsych.2020.12.025.

4. Bangasser, D.A., and Cuarenta, A. (2021). Sex differences in anxiety and depression: circuits and mechanisms. Nat Rev Neurosci 22, 674–684. 10.1038/s41583-021-00513-0.

5. Mohammadi, S., Seyedmirzaei, H., Salehi, M.A., Jahanshahi, A., Zakavi, S.S., Dehghani Firouzabadi, F., and Yousem, D.M. (2023). Brain-based Sex Differences in Depression: A Systematic Review of Neuroimaging Studies. Brain Imaging and Behavior 17, 1–29.

6. Thomas, J.T., Thorp, J.G., Huider, F., Grimes, P.Z., Wang, R., Youssef, P., Coleman, J.R.I., Byrne, E.M., Adams, M., Medland, S.E., et al. (2025). Sex-stratified genome-wide association meta-analysis of major depressive disorder. Nature Communications 16, 7960. 10.1038/s41467-025-63236-1.

7. Seney, M.L., Huo, Z., Cahill, K., French, L., Puralewski, R., Zhang, J., Logan, R.W., Tseng, G., Lewis, D.A., and Sibille, E. (2018). Opposite Molecular Signatures of Depression in Men and Women. Biological Psychiatry 84, 18–27. 10.1016/j.biopsych.2018.01.017.

8. Issler, O., and Nestler, E.J. (2018). The molecular basis for sex differences in depression susceptibility. Current Opinion in Behavioral Sciences 23, 1–6. 10.1016/j.cobeha.2017.12.019.

9. Niu, J., Verkhratsky, A., Butt, A., and Yi, C. (2025). Oligodendroglia and Myelin: Supporting the Connectome. Adv Neurobiol 43, 1–37. 10.1007/978-3-031-87919-7_1.

10. Butt, A.M., Niu, J., Yi, C., and Verkhratsky, A. Physiology of oligodendroglia. 10.1152/physrev.00023.2025.

11. Zhou, B., Zhu, Z., Ransom, B.R., and Tong, X. (2021). Oligodendrocyte lineage cells and depression. Mol Psychiatry 26, 103–117. 10.1038/s41380-020-00930-0.

12. Gan, Y., Ye, M., Wen, J., Zeng, Q., Fu, J., Yang, H., and Shi, Y. (2026). Oligodendrocyte dysfunction in major depressive disorder: Mechanistic insights and emerging therapies. Psychiatry Research 359, 117025. 10.1016/j.psychres.2026.117025.

13. Kokkosis, A.G., Madeira, M.M., Mullahy, M.R., and Tsirka, S.E. (2022). Chronic stress disrupts the homeostasis and progeny progression of oligodendroglial lineage cells, associating immune oligodendrocytes with prefrontal cortex hypomyelination. Mol Psychiatry 27, 2833–2848. 10.1038/s41380-022-01512-y.

14. Nagy, C., Maitra, M., Tanti, A., Suderman, M., Théroux, J.-F., Davoli, M.A., Perlman, K., Yerko, V., Wang, Y.C., Tripathy, S.J., et al. (2020). Single-nucleus transcriptomics of the prefrontal cortex in major depressive disorder implicates oligodendrocyte precursor cells and excitatory neurons. Nat Neurosci 23, 771–781. 10.1038/s41593-020-0621-y.

15. Ma, G., Wu, J., Wu, Y., Yang, J., Zhang, J., Huang, Y., Tian, R., Zhou, X., Tan, X., Li, Y., et al. (2025). Single-cell transcriptomics reveals Sox6 positive interneurons enriched in the prefrontal cortex of female mice vulnerable to chronic social stress. Molecular Psychiatry, 1–14. 10.1038/s41380-025-03088-9.

16. Gibson, E.M., Purger, D., Mount, C.W., Goldstein, A.K., Lin, G.L., Wood, L.S., Inema, I., Miller, S.E., Bieri, G., Zuchero, J.B., et al. (2014). Neuronal Activity Promotes Oligodendrogenesis and Adaptive Myelination in the Mammalian Brain. Science 344, 1252304. 10.1126/science.1252304.

17. Issler, O., van der Zee, Y.Y., Ramakrishnan, A., Xia, S., Zinsmaier, A.K., Tan, C., Li, W., Browne, C.J., Walker, D.M., Salery, M., et al. (2022). The long noncoding RNA FEDORA is a cell type– and sex-specific regulator of depression. Sci Adv 8, eabn9494. 10.1126/sciadv.abn9494.

18. Kato, D. (2025). Impact of Sex Differences in Oligodendrocytes and Their Progenitor Cells on the Pathophysiology of Neuropsychiatric Disorders. J Nippon Med Sch 92, 226–233. 10.1272/jnms.JNMS.2025_92-306.

19. Su, J., Song, Y., Zhu, Z., Huang, X., Fan, J., Qiao, J., and Mao, F. (2024). Cell–cell communication: new insights and clinical implications. Sig Transduct Target Ther 9, 1–52. 10.1038/s41392-024-01888-z.

20. Jin, S., and Ramos, R. (2022). Computational exploration of cellular communication in skin from emerging single-cell and spatial transcriptomic data. Biochemical Society Transactions 50, 297–308. 10.1042/BST20210863.

21. Maitra, M., Mitsuhashi, H., Rahimian, R., Chawla, A., Yang, J., Fiori, L.M., Davoli, M.A., Perlman, K., Aouabed, Z., Mash, D.C., et al. (2023). Cell type specific transcriptomic differences in depression show similar patterns between males and females but implicate distinct cell types and genes. Nature Communications 14, 2912. 10.1038/s41467-023-38530-5.

22. Jin, S., Guerrero-Juarez, C.F., Zhang, L., Chang, I., Ramos, R., Kuan, C.-H., Myung, P., Plikus, M.V., and Nie, Q. (2021). Inference and analysis of cell-cell communication using CellChat. Nat Commun 12, 1088. 10.1038/s41467-021-21246-9.

23. Al Shweiki, M.R., Oeckl, P., Steinacker, P., Barschke, P., Dorner-Ciossek, C., Hengerer, B., Schönfeldt-Lecuona, C., and Otto, M. (2020). Proteomic analysis reveals a biosignature of decreased synaptic protein in cerebrospinal fluid of major depressive disorder. Transl Psychiatry 10, 144. 10.1038/s41398-020-0825-7.

24. Zhao, W., Johnston, K.G., Ren, H., Xu, X., and Nie, Q. (2023). Inferring neuron-neuron communications from single-cell transcriptomics through NeuronChat. Nature Communications 14, 1128. 10.1038/s41467-023-36800-w.

25. Piñero, J., Bravo, À., Queralt-Rosinach, N., Gutiérrez-Sacristán, A., Deu-Pons, J., Centeno, E., García-García, J., Sanz, F., and Furlong, L.I. (2017). DisGeNET: a comprehensive platform integrating information on human disease-associated genes and variants. Nucleic Acids Res 45, D833–D839. 10.1093/nar/gkw943.

26. Kamran, M., Bibi, F., ur. Rehman, Asim., and Morris, D.W. (2022). Major Depressive Disorder: Existing Hypotheses about Pathophysiological Mechanisms and New Genetic Findings. Genes (Basel) 13, 646. 10.3390/genes13040646.

27. Pitsillou, E., Bresnehan, S.M., Kagarakis, E.A., Wijoyo, S.J., Liang, J., Hung, A., and Karagiannis, T.C. (2020). The cellular and molecular basis of major depressive disorder: towards a unified model for understanding clinical depression. Mol Biol Rep 47, 753–770. 10.1007/s11033-019-05129-3.

28. Wang, Y., Xiong, Z., Chen, M., and Zhu, Z. (2026). Myelin damage in major depressive disorder: Insights from neuroimaging, molecular mechanisms, and therapeutic perspectives. Neuroscience & Biobehavioral Reviews 184, 106607. 10.1016/j.neubiorev.2026.106607.

29. Morabito, S., Reese, F., Rahimzadeh, N., Miyoshi, E., and Swarup, V. (2023). hdWGCNA identifies co-expression networks in high-dimensional transcriptomics data. Cell Rep Methods 3, 100498. 10.1016/j.crmeth.2023.100498.

30. Rong, Y., Yan, W., Gao, Z., Yang, Y., Xu, C., and Zhang, C. (2025). NRXN3-NLGN1 complex influences the development of depression induced by maternal separation in rats. Brain Research 1858, 149659. 10.1016/j.brainres.2025.149659.

31. Wang, J., Sudhof, T., and Wernig, M. (2025). Distinct mechanisms control the specific synaptic functions of Neuroligin 1 and Neuroligin 2. EMBO Rep 26, 860–879. 10.1038/s44319-024-00286-4.

32. Baer, K., Essrich, C., Benson, J.A., Benke, D., Bluethmann, H., Fritschy, J.-M., and Lüscher, B. (1999). Postsynaptic clustering of γ-aminobutyric acid type A receptors by the γ3 subunit in vivo. Proceedings of the National Academy of Sciences 96, 12860–12865. 10.1073/pnas.96.22.12860.

33. Gomez, A.M., Traunmüller, L., and Scheiffele, P. (2021). Neurexins: molecular codes for shaping neuronal synapses. Nat Rev Neurosci 22, 137–151. 10.1038/s41583-020-00415-7.

34. Yao, Z., van Velthoven, C.T.J., Kunst, M., Zhang, M., McMillen, D., Lee, C., Jung, W., Goldy, J., Abdelhak, A., Aitken, M., et al. (2023). A high-resolution transcriptomic and spatial atlas of cell types in the whole mouse brain. Nature 624, 317–332. 10.1038/s41586-023-06812-z.

35. Young, K.M., Psachoulia, K., Tripathi, R.B., Dunn, S.-J., Cossell, L., Attwell, D., Tohyama, K., and Richardson, W.D. (2013). Oligodendrocyte Dynamics in the Healthy Adult CNS: Evidence for Myelin Remodeling. Neuron 77, 873–885. 10.1016/j.neuron.2013.01.006.

36. Wang, X., Zeng, C., Wu, Z., Lu, M., Wang, X., Xiu, Y., Wang, Q., Wang, S., Chen, X., Shen, Y., et al. (2025). Chromatin remodeling factor BAF155 coordinates oligodendroglial-neuronal communications linked to regional myelination and autism-like behavioral deficits in mice. Nat Commun 17, 1165. 10.1038/s41467-025-67930-y.

37. Yamazaki, R., Azuma, M., Osanai, Y., Kouki, T., Inagaki, T., Kakita, A., Takao, M., and Ohno, N. (2025). Type I collagen secreted in white matter lesions inhibits remyelination and functional recovery. Cell Death Dis 16, 285. 10.1038/s41419-025-07633-w.

38. Nicoletti, A., Baschi, R., Cicero, C.E., Iacono, S., Re, V.L., Luca, A., Schirò, G., and Monastero, R. (2023). Sex and gender differences in Alzheimer’s disease, Parkinson’s disease, and Amyotrophic Lateral Sclerosis: A narrative review. Mechanisms of Ageing and Development 212, 111821. 10.1016/j.mad.2023.111821.

39. Rainville, J.R., Lipuma, T., and Hodes, G.E. (2022). Translating the Transcriptome: Sex Differences in the Mechanisms of Depression and Stress, Revisited. Biol Psychiatry 91, 25–35. 10.1016/j.biopsych.2021.02.003.

40. Medina, A., Watson, S.J., Bunney, W., Myers, Richard. M., Schatzberg, A., Barchas, J., Akil, H., and Thompson, R.C. (2016). Evidence for alterations of the glial syncytial function in Major Depressive Disorder. J Psychiatr Res 72, 15–21. 10.1016/j.jpsychires.2015.10.010.

41. Sild, M., Ruthazer, E.S., and Booij, L. (2017). Major depressive disorder and anxiety disorders from the glial perspective: Etiological mechanisms, intervention and monitoring. Neuroscience & Biobehavioral Reviews 83, 474–488. 10.1016/j.neubiorev.2017.09.014.

42. Lin, S.-S., Zhou, B., Liu, S.-L., Ren, X.-Y., Guo, J., Tong, J.-L., Chen, B.-J., Jiang, R.-T., Semyanov, A., Yi, C., et al. (2025). Astrocyte Ezrin defines resilience to stress-induced depressive behaviours in mice. Natl Sci Rev 13, nwaf480. 10.1093/nsr/nwaf480.

43. Lin, S.-S., Zhou, B., Chen, B.-J., Jiang, R.-T., Li, B., Illes, P., Semyanov, A., Tang, Y., and Verkhratsky, A. (2023). Electroacupuncture prevents astrocyte atrophy to alleviate depression. Cell Death Dis 14, 343. 10.1038/s41419-023-05839-4.

44. Sanadgol, N., Miraki Feriz, A., Lisboa, S.F., and Joca, S.R.L. (2023). Putative role of glial cells in treatment resistance depression: An updated critical literation review and evaluation of single-nuclei transcriptomics data. Life Sciences 331, 122025. 10.1016/j.lfs.2023.122025.

45. Verkhratsky, A., Lee, C.J., Chun, H., Göritz, C., Harkany, T., Lee, J.-H., Lee, S., Lindskog, M., Koh, W., Mulder, J., et al. (2026). Curing the brain: in search for new astrocyte-specific therapies. Exp Mol Med 58, 1086–1127. 10.1038/s12276-026-01712-4.

46. Ma, Z., Zhang, W., Wang, C., Su, Y., Yi, C., and Niu, J. (2024). A New Acquaintance of Oligodendrocyte Precursor Cells in the Central Nervous System. Neurosci Bull 40, 1573–1589. 10.1007/s12264-024-01261-8.

47. Yi, C., Verkhratsky, A., and Niu, J. (2023). Pathological potential of oligodendrocyte precursor cells: terra incognita. Trends in Neurosciences 46, 581–596. 10.1016/j.tins.2023.04.003.

48. Pantazatos, S.P., Huang, Y., Rosoklija, G.B., Dwork, A.J., Arango, V., and Mann, J.J. (2017). Whole-transcriptome brain expression and exon-usage profiling in major depression and suicide: evidence for altered glial, endothelial and ATPase activity. Mol Psychiatry 22, 760–773. 10.1038/mp.2016.130.

49. Birey, F., Kloc, M., Chavali, M., Hussein, I., Wilson, M., Christoffel, D.J., Chen, T., Frohman, M.A., Robinson, J.K., Russo, S.J., et al. (2015). Genetic and Stress-Induced Loss of NG2 Glia Triggers Emergence of Depressive-like Behaviors through Reduced Secretion of FGF2. Neuron 88, 941–956. 10.1016/j.neuron.2015.10.046.

50. Wang, Y., Su, Y., Yu, G., Wang, X., Chen, X., Yu, B., Cheng, Y., Li, R., Sáez, J.C., Yi, C., et al. (2021). Reduced Oligodendrocyte Precursor Cell Impairs Astrocytic Development in Early Life Stress. Adv Sci (Weinh) 8, 2101181. 10.1002/advs.202101181.

51. DeCasien, A.R., Auluck, P., Liu, S., Feng, N., Elkahloun, A.G., Xu, Q., Marenco, S., Cookson, M.R., and Raznahan, A. (2026). Sex effects on gene expression across the human cerebral cortex at cell type resolution. Science 392, eaea9063. 10.1126/science.aea9063.

52. Breton, J.M., Barraza, M., Hu, K.Y., Frias, S.J., Long, K.L.P., and Kaufer, D. (2021). Juvenile exposure to c leads to long-lasting alterations in grey matter myelination in adult female but not male rats. Neurobiol Stress 14, 100319. 10.1016/j.ynstr.2021.100319.

53. Jastrzębska, J., Frankowska, M., Wesołowska, J., Filip, M., and Smaga, I. (2025). Dietary Intervention with Omega-3 Fatty Acids Mitigates Maternal High-Fat Diet-Induced Behavioral and Myelin-Related Alterations in Adult Offspring. Curr Neuropharmacol 23, 329–348. 10.2174/1570159X23666241014164940.

54. Ho, T.C., Sisk, L.M., Kulla, A., Teresi, G.I., Hansen, M.M., Wu, H., and Gotlib, I.H. (2021). Sex differences in myelin content of white matter tracts in adolescents with depression. Neuropsychopharmacology 46, 2295–2303. 10.1038/s41386-021-01078-3.

55. Li, J., and Monk, K.R. (2024). Synapses shape oligodendrocyte precursor cell development and predict myelination location. Nat Neurosci 27, 217–218. 10.1038/s41593-023-01555-6.

56. Silva, R.H., Pedro, L.C., Manosso, L.M., Gonçalves, C.L., and Réus, G.Z. (2024). Pre-and Post-Synaptic protein in the major depressive Disorder: From neurobiology to therapeutic targets. Neuroscience 556, 14–24. 10.1016/j.neuroscience.2024.07.050.

57. Bang, M.L., and Owczarek, S. (2013). A Matter of Balance: Role of Neurexin and Neuroligin at the Synapse. Neurochem Res 38, 1174–1189. 10.1007/s11064-013-1029-9.

58. Seeker, L.A., Bestard-Cuche, N., Jäkel, S., Kazakou, N.-L., Bøstrand, S.M.K., Wagstaff, L.J., Cholewa-Waclaw, J., Kilpatrick, A.M., Van Bruggen, D., Kabbe, M., et al. (2023). Brain matters: unveiling the distinct contributions of region, age, and sex to glia diversity and CNS function. Acta Neuropathol Commun 11, 84. 10.1186/s40478-023-01568-z.

59. Irene, S.-S., Borja, G.-C., Rubén, G.-R., Cristina, G.-R., Lucas, B.-M., Héctor, C., María, de la I.-V., Sara, G.-P., Vanja, T., R, H.M., et al. (2024). Single cell landscape of sex differences in the progression of multiple sclerosis. Preprint at Research Square, 10.21203/rs.3.rs-5482526/v1 10.21203/rs.3.rs-5482526/v1.

60. Poulopoulos, A., Aramuni, G., Meyer, G., Soykan, T., Hoon, M., Papadopoulos, T., Zhang, M., Paarmann, I., Fuchs, C., Harvey, K., et al. (2009). Neuroligin 2 drives postsynaptic assembly at perisomatic inhibitory synapses through gephyrin and collybistin. Neuron 63, 628–642. 10.1016/j.neuron.2009.08.023.

61. Bai, X., Kirchhoff, F., and Scheller, A. (2021). Oligodendroglial GABAergic Signaling: More Than Inhibition! Neurosci Bull 37, 1039–1050. 10.1007/s12264-021-00693-w.

62. Hamilton, N.B., Clarke, L.E., Arancibia-Carcamo, I.L., Kougioumtzidou, E., Matthey, M., Káradóttir, R., Whiteley, L., Bergersen, L.H., Richardson, W.D., and Attwell, D. (2017). Endogenous GABA controls oligodendrocyte lineage cell number, myelination, and CNS internode length. Glia 65, 309–321. 10.1002/glia.23093.

63. Serrano-Regal, M.P., Luengas-Escuza, I., Bayón-Cordero, L., Ibarra-Aizpurua, N., Alberdi, E., Pérez-Samartín, A., Matute, C., and Sánchez-Gómez, M.V. (2020). Oligodendrocyte Differentiation and Myelination Is Potentiated via GABAB Receptor Activation. Neuroscience 439, 163–180. 10.1016/j.neuroscience.2019.07.014.

64. Cortes, L.R., Sturgeon, H., and Forger, N.G. (2023). Sexual differentiation of estrogen receptor alpha subpopulations in the ventromedial nucleus of the hypothalamus. Horm Behav 151, 105348. 10.1016/j.yhbeh.2023.105348.

65. Gaesser, J.M., and Fyffe-Maricich, S.L. (2016). Intracellular Signaling Pathway Regulation of Myelination and Remyelination in the CNS. Exp Neurol 283, 501–511. 10.1016/j.expneurol.2016.03.008.

66. Ishii, A., Furusho, M., Dupree, J.L., and Bansal, R. (2014). Role of ERK1/2 MAPK Signaling in the Maintenance of Myelin and Axonal Integrity in the Adult CNS. J Neurosci 34, 16031–16045. 10.1523/JNEUROSCI.3360-14.2014.

67. Bell-Horner, C.L., Dohi, A., Nguyen, Q., Dillon, G.H., and Singh, M. (2006). ERK/MAPK pathway regulates GABAA receptors. Journal of Neurobiology 66, 1467–1474. 10.1002/neu.20327.

68. Hernández-Hernández, E., Ledesma-Corvi, S., Yáñez-Gómez, F., Mateu-Mercader, N., Bagán, A., Escolano, C., and García-Fuster, M.J. (2026). Sex differences in the antidepressant-like response induced by the imidazoline-2 receptor compound 12d, a (3-phenylcarbamoyl-3,4-dihydro-2H-pyrrol-2-yl)phosphonate. Pharmacol Rep 78, 341–349. 10.1007/s43440-025-00810-w.

69. Negrini-Ferrari, S.E., Martínez-Martel, I., and Pol, O. (2026). Molecular Hydrogen Reverses Nociplastic Pain and Depressive-like Behaviors via Region- and Sex-Dependent Central Mechanisms. Int J Mol Sci 27, 3051. 10.3390/ijms27073051.

70. Torous, J., Linardon, J., Goldberg, S.B., Sun, S., Bell, I., Nicholas, J., Hassan, L., Hua, Y., Milton, A., and Firth, J. (2025). The evolving field of digital mental health: current evidence and implementation issues for smartphone apps, generative artificial intelligence, and virtual reality. World Psychiatry 24, 156–174. 10.1002/wps.21299.

