## Supplementary material for "Aberrant neuron-OPC synaptic transmission and deficient myelination define sex bias in major depressive disorder": Supplemenal information

### Methods

#### Acquisition of the human single-nucleus RNA-sequencing dataset

Single-nucleus RNA sequencing (snRNA-seq) datasets of the dorsolateral prefrontal cortex (dlPFC) from human MDD patients and healthy controls were acquired from NCBI GEO database under accession numbers GSE144136 and GSE213982 [1,2], composed of 37 MDD patients (20 females, 17 males) and 34 matched healthy controls (18 females, 16 males). The rigorously validated dimensionality reduction coordinates (UMAP) and the hierarchical cell-type annotations provided by the original study were adopted, ensuring maximal consistency with established neuroanatomical definitions. The integrated datasets comprised 7 broad brain cell types and 41 refined cell clusters of excitatory (20 refined clusters) and inhibitory neurons (10 refined clusters), OLs (3 clusters), astrocytes (2 clusters), OPCs (3 clusters), endothelial cells (1 cluster), and microglia (1 cluster), including 1 unassigned cluster denoted as Mix. The normalized gene expression matrices, coupled with the imported metadata, served as the foundational input for all subsequent downstream interaction and lineage-specific analyses.

#### Inference of intercellular communication networks

To infer intercellular communication networks, we utilized the R packages CellChat [3] (version 1.1.3) and NeuronChat [4] (version 1.0.0). The identification of signaling pathways and specific interactions and the comparison analyses between MDD and control groups followed the CellChat tutorial on comparing multiple datasets. Customized CellChat functions of netAnalysis_signalingRole_scatter, netAnalysis_signalingChanges_scatter, and netAnalysis_contribution were ultilized for displaying sender-receiver roles of cell types, disease-associated signaling pathway alterations and quantifying contribution of each ligand-receptor pair with enhanced visual representation. We additionally employed NeuronChat, a cell-cell communication inference tool advantaged by its incorporation of neurotransmitter signaling to quantify the interaction landscape within the oligodendroglial lineage cells in MDD pathogenesis. Group-wise comparisons within NeuronChat were conducted following an analogous workflow.

#### Sex-stratified network comparison and topological analysis

#### Quantification of global network attenuation

The overall network attenuations were quantitatively assessed for both female and male cohorts independently. For each cohort, the global interaction strength was mathematically defined as the sum of the absolute communication probabilities across the entire inferred intercellular network matrix. Subsequently, the magnitude of network attenuation in MDD was calculated as the percentage reduction in global interaction strength relative to its sex-matched control cohort, utilizing the following formula:

$$Attenuation\ (\%) = (1 - \frac{\text{Strength}_{MDD}}{Strength_{Control}}) \times 100$$

Statistical significance of the observed network attenuation was determined via a permutation test as detailed in the Permutation analysis.

**GWAS integration and cell-type-specific risk scoring**

The largest sex-stratified genome-wide association study (GWAS) for human MDD to date (comprising a total sample size of: 130,471 female and 64,805 male MDD patients, 159,521 female and 132,185 male controls [5]) was integrated. The sex-specific MDD risk genes derived from this comprehensive GWAS were mapped onto our previously acquired human snRNA-seq dataset to assess the genetic associations across cell types. To quantify this cellular risk burden, we employed the AddModuleScore function in the Seurat [6] R package (version 5.1.0), a robust method for calculating single-cell gene set enrichment scores. Notably, this scoring methodology was consistently applied for all subsequent single-cell gene set enrichment quantifications throughout the study.

**Rank-shift analysis of communication hubs**

Network centrality was performed by quantifying the total outgoing and incoming signaling strengths, contributing the top 30% of cell clusters as communication hubs. Sex-stratified rank-shift analyses were conducted on intersecting hubs preserved across conditions and visualized using 2D scatter plots to assess broadcasting fidelity and receiver stability under pathological stress.

**Spatial visualization and distance-based ranking of cellular perturbations**

The aggregated outgoing and incoming interaction strengths of each cell cluster across four cohorts were projected onto 2D scatter plots, and three spatial distance metrics were computed to reflect their distinct extent of pathological deviation. Euclidean distance between their healthy and disease coordinates was computed to delineate the global perturbations, while the horizontal (x-axis) and vertical (y-axis) displacements were calculated to capture signaling alterations in transmission and reception. In line with that, distance rankings were performed for each sex to exhibit cellular perturbations in sex-specific MDD pathogenesis.

**Identification of perturbed signaling pathways**

We computed the differential interaction strength for each signaling pathway by comparing MDD versus control states within each sex, based on the values derived from the netP slot in CellChat. These interaction strength changes were ranked in descending order by their absolute magnitude and visualized in an elbow plot. The inflection point at a value of 5 was utilized as the threshold for identifying significant pathway perturbations in both sexes. The statistical significance was confirmed using the permutation test as detailed in the Permutation analysis.

**Pathological and functional interpretation of oligodendroglia**

### Subclustering

The oligodendroglial lineage cells were computationally isolated and subjected to high-resolution subclustering in the Seurat [6] R package (version 5.1.0). Based on the Harmony-corrected PCA components, this iterative clustering approach identified three primary functional subsets (Ident 1, Ident 2, and Ident 3), characterized by distinct transcriptomic profiles and canonical marker expression. Furthermore, the cells of Ident 3 were extracted for secondary refined subclustering, yielding three subclusters through an analogous workflow.

### Cell proportion fold change analysis

The disease-associated shift in the cellular proportion of oligodendroglial subsets was quantified as a normalized fold change. For each subset, the proportion of each cohort was calculated as the number of cells in that subset divided by the total cells of the cohort. The fold change for an MDD cohort was then defined as its proportion divided by the corresponding sex-matched control, with the latter normalized to 1.

### DisGeNET analysis

With the list of ident-specific top 150 transcriptomic signatures, we ran gene2disease from R package disgenet2r (version 1.2.2) with database = "CURATED" and other parameters set to default to find the neuropsychiatric disorders for which our ident-specific signatures showed an enrichment reported in DisGeNET [7]. The disease relevance with the representative neuropsychiatric disorders of MDD, bipolar disorder, schizophrenia, autistic disorder and alcohol abuse was compared among the subsets.

### Functional enrichment analysis

Gene Set Enrichment Analysis (GSEA [8]) and Gene Ontology (GO) enrichment analyses were performed on the differentially expressed genes via gseGO and enrichGO functions in R package clusterProfiler (version 4.14.6), with a $p$-value < 0.05 deemed statistically significant. R packages of GseaVis (version 0.1.1) and ClusterGVis (version 0.1.4) for enhanced visualization were also utilized.

### Pseudotime trajectory analysis

Pseudotime trajectory inference was performed with the oligodendroglial Ident3 using the R package Slingshot [9] (version 2.14.0). This delineated a unified trajectory mapping the oligodendroglial differentiation of Ident3, with the male and female datasets combined due to the limitation of the number of cells.

### High-dimensional weighted gene co-expression network analysis (hdWGCNA)

High-dimensional weighted gene co-expression network analysis (hdWGCNA [10]) was performed to identify co-expression modules using snRNA-seq data. Specifically, module preservation analysis with hdWGCNA, the statistical framework that ensuring the evaluation of the degree to which co-expression modules identified in control dataset are "preserved" in MDD dataset was conducted using R package hdWGCNA (version 0.4.1). The preservation analyses followed the hdWGCNA tutorial on identifying and projecting modules and visualizing the preservation stats.

**RNA-sequencing and analysis**

OPCs were isolated by immunopanning, and RNA was extracted from the isolated cells by Trizol (Thermo, Cat. No. 15596026) according to the manufacturer's protocol. RNA-seq and subsequent upstream analysis were performed by BGI-Shenzhen, China. The GRCm38 was used for genome reference. Essentially, differential expression analysis was performed using the DESeq2 (v1.44.0). Genes with FDR-adjusted *p*-value below 0.05 were considered statistically significant. For region-specific visualization of differentially expressed genes, an additional filter of absolute $\log_2$fold-change above 1 (equivalent to 2-fold change) was applied. To identify concordant pathway dysregulations across the mPFC and hippocampus, functional enrichment analysis was performed on the statistically significant genes showing consistent directional changes in both regions, without imposing the fold-change filter. Downstream analysis and data visualization were conducted using R packages of clusterProfiler (version 4.14.6), ComplexHeatmap (version 2.20.0) and enrichplot (version 1.26.6).

**Gene expression analysis from online mouse scRNA-seq platform**

The Mouse Whole-Brain Transcriptomic Cell Type Atlas was accessed through the Allen Institute for Brain Science portal (https://portal.brain-map.org) [11]. This dataset includes single-cell RNA-seq data from approximately 4 million cells, encompassing diverse cell types across the entire mouse brain. To explore the expression patterns of the target genes and various Gabr subtypes, we utilized the portal's search and interactive visualization tools. The platform offers UMAP plots and heatmaps illustrating the expression of *Gabrg3*, *Nrxn3*, *Nlgn1* and *Gephyrin* across different cell types. Ridgeline plots were generated using the R package ggridges (version 0.5.7) based on data retrieved from the same atlas.

**Mice**

All animal studies were performed under the guidelines of laboratory animal welfare and ethics committee of the Third Military Medical University (AMUWEC20223048). All mice were housed in a temperature- and humidity-controlled environments with free access to standard chow and water and on a 12 h/12 h light/dark cycle.

**PDGFRα$^{\mathrm{CreER}}$; Gabrg3$^{\mathrm{fl/fl}}$**

*PDGFRα*$^{\mathrm{CreER}}$ mice were acquired from Dr. Stephen Fancy at the University of California, San Francisco [12]. Gabrg3-flox mice were generated by Gempharmatech Co.,Ltd. (China). The mice were then crossed with *PDGFRα*$^{\mathrm{CreER}}$ mice to generate *PDGFRα*$^{\mathrm{CreER}}$; *Gabrg*3$^{\mathrm{fl/fl}}$ conditional KO mice. Non-CreER; *Gabrg3*$^{\mathrm{fl/fl}}$ littermates were used as the control group. Mice were administered tamoxifen (10 mg/kg/d, gavage) for 5 consecutive days from P4 to P8 to induce cre-mediated reorganization.

**NG2Cre$^{\mathrm{ERT}}$; Tau-mGFP**

The NG2$^{CreERT}$ line (The Jackson Laboratory, Catalog # 008538) was crossed with the Tau-mGFP (The Jackson Laboratory, Catalog # 021162) line to obtain NG2$^{CreERT}$; Tau-mGFP mice. Mice were administered tamoxifen (10 mg/kg/day, gavage) for 5 consecutive days to induce Cre-mediated recombination. Upon differentiation of targeted NG2 positive OPCs and the subsequent onset of Tau expression, mGFP is expressed in line with Tau.

**Behavior tests**

Mice were handled daily for five days before the test. The apparatus was wiped with 80% alcohol between each trial. All tests were conducted between 9 am and 6 pm. The experimenters were blinded for the grouping. VisuTrack Animal Behavior Analysis Software (Shanghai XinRuan) was used for data collection and analysis.

**Tail suspension test**

Depression-related behavior was evaluated using the tail suspension test. Each mouse was suspended by securing its tail with adhesive tape to the top of a testing chamber (55 × 15 × 11.5 cm), ensuring the head was positioned approximately 25 cm above the chamber floor. The immobility duration was recorded over a 10-minute period.

**Forced swimming test**

The forced swimming test served to assess depression-like behavior. Mice were individually placed into a cylindrical arena (10 × 25 cm) filled with water to a depth of 15 cm at 25°C. The duration of immobility was quantified during a 6-minute session.

**Elevated plus maze**

Anxiety-like behavior was measured using the elevated plus maze. Each mouse was placed at the intersection of the open and closed arms, facing an open arm, and allowed to explore the maze for 10 minutes. Anxiety levels were determined by comparing the time spent or distance traveled in the closed arms.

**Open field test**

Locomotor function and anxiety-like behavior were assessed via the open field test. Mice were placed in the center of a 50 × 50 cm arena and permitted to explore freely for 15 minutes. Total distance traveled, as well as time spent and distance traveled in the central zone, were recorded. A lower time or distance in the center indicated heightened anxiety-like behavior. Data acquisition and analysis were performed using VisuTrack Animal Behavior Analysis Software.

**The three-chamber test**

Sociability and preference for social novelty were evaluated using the three-chamber test. The apparatus comprised left, right, and center chambers. Test mice were first allowed to freely explore all three chambers for 10 minutes. In the sociability phase (labeled Stranger1-Object, S1-O in figures), an unfamiliar same-sex, age-matched mouse was placed into a cage in either the left or right chamber, while an empty cage was placed in the opposite chamber. Sociability was measured by recording sniffing time directed toward the caged mouse (Stranger1) versus the empty cage. In the subsequent social novelty preference phase (labeled Stranger1-Stranger2, S1-S2 in figures), a second unfamiliar mouse was placed into the previously empty cage, and sniffing time toward the novel mouse (Stranger2) was compared with that toward the familiar mouse (Stranger1).

**The self-grooming test**

This test assessed repetitive behaviors in mice. Each mouse was placed in a transparent chamber and given 10 minutes to explore and acclimate freely, followed by another 10 minutes of recording. The total time spent on self-grooming behavior was then calculated.

**Novel object recognition test**

This test evaluated short-term memory function. For two days prior to the experiment, mice were placed daily for 10 minutes in a 25 × 25 × 40 cm chamber to habituate to the environment. During the trial, two identical objects were placed in the chamber, and each mouse was allowed to explore them for 5 minutes. Two hours later, one of the objects was replaced with a novel one differing in shape and color, and the mouse was again allowed to explore for 5 minutes. The time spent exploring the novel object versus the familiar one was recorded.

**Real time-PCR**

Total RNA was extracted using RNeasy Plus Mini Kit (Qiagen, Cat# 74134). The PrimeScript RT Reagent Kit (Takara) was used for reverse transcription. The Accurate 96 Real Time PCR System (DLAB) and FastStart Universal SYBR Green Master Mix (Roche, 04913850001) were used for the qPCR experiment.

**Immunohistochemistry**

Mice were intracardially perfused with 4% PFA, and brains were removed and fixed in 4% PFA overnight at 4°C. Then brains were cryoprotected with 30% sucrose for 3 days before cryosection at 20 μm. Sections were blocked with 5% bovine serum albumin (BSA) with 0.25% Triton-X 100 for two hours at room temperature before incubated with primary antibodies overnight at 4°C, followed by secondary antibodies for one hour at room temperature. The primary antibodies include: Goat anti-PDGFR-α (1:200, R&D, AF1062); Rat anti-GFP (1:200, Abcam, AB5450); Rabbit anti-Gabrg3 (1:200, Abbexa, abx032311); Rabbit anti-Gephyrin (1:200, Proteintech, 12681-1-AP); Rabbit anti-NRXN3 (1:200, Invitrogin, PA5-37189); Mouse anti-NLGN1 (1:200, Proteintech, 66964-1-Ig); Mouse anti-IP3RII (1:100, Santa Cruz, sc398434); Rabbit anti-SOX10 (1:200, CST, 89356T); Guinea anti-vGlut1 (1:2000, Synaptic Systems, 135304); Mouse anti-vGAT (1:1000, Synaptic Systems, 131011); Rabbit anti-PV (1:500, Abacam, ab11427); Rat anti-MBP (1:500, Millipore, MAB395). The Olympus VS200 Research

Slide Scanner (Olympus) and Ixplore SpinSR confocal microscope (Olympus) were used for imaging. Fluorescent images were analyzed using the CellSens (Olympus) and ImageJ (NIH, USA).

**Electron microscopy**

Preparation of electron microscopy of the optic nerves was performed as previously described [13]. Mice were anesthetized and perfused intracardially with 0.1 M PB, followed by 2.5% glutaraldehyde and 4% glutaraldehyde in 0.1 M PB perfusion and post-fixed overnight. Then, optic nerves were isolated and stained with osmium tetroxide overnight and were subjected to a series of ethanol dehydration treatments. The tissue embedding was performed in TAAB resin to acquire 1 μm sections. More than 200 axons were examined using electron microscopy. The myelinated axon numbers and G-ratios were calculated to assess the demyelination and remyelination.

**Calcium imaging**

As described previously [13], acute brain slices and cultured OPCs were incubated with calcium probe Rhod-5 at 37°C in the dark for 30 min. Confocal images were captured by Ixplore SpinSR confocal microscope (Olympus) at 200ms per frame and analyzed by the ImageJ software.

**Electrophysiology**

The N-methyl-D-glucamine protective recovery method was used to prepare brain slices [14]. NMDG-HEPES aCSF contains (in mM): 92 NMDG, 2.5 KCl, 1.25 NaH2PO4, 30 NaHCO3, 20 HEPES, 25 glucose, 2 thiourea, 5 Na-ascorbate,3 Na-pyruvate, 0.5 CaCl2·2H2O, and 10 MgSO4·7H2O. HEPES with aCSF contains (in mM): 92 NaCl, 2.5 KCl, 1.25 NaH2PO4, 30 NaHCO3, 20 HEPES, 25 glucose, 2 thiourea, 5 Na-ascorbate, 3Na-pyruvate, 2 CaCl2·2H2O, and 2 MgSO4·7H2O. For spontaneous excitatory postsynaptic currents (sEPSCs) and inhibitory postsynaptic currents (sIPSCs) recording, whole-cell patch-clamp recordings were performed using borosilicate glass pipettes. sEPSCs were recorded at a holding potential Of -70 mV and slPSCs were recorded at a holding potential of + 10 mV in regular ACSF. A MultiClamp 700B amplifier and pCLAMP10 software were used for electrophysiology (Axon Instruments). Minianalysis software was used for data analysis.

**Statistics**

**Permutation analysis**

To rigorously validate the statistical significance of the observed network alterations, a label-shuffling permutation analysis was implemented. The diagnostic labels of MDD patients versus controls were randomly permuted 2000 times, within the female and male datasets separately. Downstream network inferences (including both CellChat and NeuronChat analyses) were re-run for each permutation. For evaluating the reduction in global interaction strength and the identification of significantly perturbed signaling pathways, the *p*-value was defined as the proportion of permutations yielding a value greater than or equal to the actual observed metric. For network features encompassing the top 10 most divergent cell clusters, the magnitude of dysregulation of NRXN pathway and its core ligand-receptor pairs, and the oligodendroglial-specific alteration profiles derived from NeuronChat, the $p$-value was determined by two-sided Kolmogorov-Smirnov test in GraphPad Prism based on the distributions constructed by the 2,000 permutations.

#### Quantification of cell numbers

Immunostaining-positive cells with visible somata were manually counted, and the counts were normalized to the area of the region of interest.

#### Quantification of fluorescence intensity or positive area

Fluorescence intensity and the percentage of positive area were analyzed using Fiji software. A threshold was applied to identify positive fluorescence signals, after which the intensity and proportional area within the region of interest were measured. The same threshold was used across experimental groups within each batch.

#### Quantification of synaptic element immunostaining in OPCs

Analysis was performed using Imaris 9.0 following a previously described method [15]. Briefly, OPC surfaces were reconstructed based on PDGFRα signals, which were then used to mask synaptic element staining. Surface rendering was applied to the masked synaptic signals, and the number of objects was quantified.

#### Statistical analysis

GraphPad Prism 9.0 (GraphPad Software, San Diego, CA, USA) was used to determine statistical significance. Data are presented as mean ± standard error of the mean (SEM). Comparisons between two experimental groups were made using unpaired two-tailed t-tests. P-values < 0.05 were considered statistically significant, with significance levels indicated as: *$p < 0.05$, **$p < 0.01$, ***$p < 0.001$, ****$p < 0.0001$. Data distribution was assumed to be normal, although this was not formally tested. Sample sizes were not predetermined by statistical methods but were comparable to those reported in previous studies. Each experiment was repeated at least three times.

### Reference


1. Maitra, M., Mitsuhashi, H., Rahimian, R., Chawla, A., Yang, J., Fiori, L.M., Davoli, M.A., Perlman, K., Aouabed, Z., Mash, D.C., et al. (2023). Cell type specific transcriptomic differences in depression show similar patterns between males and females but implicate distinct cell types and genes. Nature Communications *14*, 2912. https://doi.org/10.1038/s41467-023-38530-5.

2. Nagy, C., Maitra, M., Tanti, A., Suderman, M., Théroux, J.-F., Davoli, M.A., Perlman, K., Yerko, V., Wang, Y.C., Tripathy, S.J., et al. (2020). Single-nucleus transcriptomics of the prefrontal cortex in major depressive disorder implicates oligodendrocyte precursor cells and excitatory neurons. Nat Neurosci *23*, 771–781. https://doi.org/10.1038/s41593-020-0621-y.

3. Jin, S., Guerrero-Juarez, C.F., Zhang, L., Chang, I., Ramos, R., Kuan, C.-H., Myung, P., Plikus, M.V., and Nie, Q. (2021). Inference and analysis of cell-cell communication using CellChat. Nat Commun *12*, 1088. https://doi.org/10.1038/s41467-021-21246-9.

4. Zhao, W., Johnston, K.G., Ren, H., Xu, X., and Nie, Q. (2023). Inferring neuron-neuron communications from single-cell transcriptomics through NeuronChat. Nature Communications *14*, 1128. https://doi.org/10.1038/s41467-023-36800-w.

5. Thomas, J.T., Thorp, J.G., Huider, F., Grimes, P.Z., Wang, R., Youssef, P., Coleman, J.R.I., Byrne, E.M., Adams, M., Medland, S.E., et al. (2025). Sex-stratified genome-wide association meta-analysis of major depressive disorder. Nature Communications *16*, 7960. https://doi.org/10.1038/s41467-025-63236-1.

6. Hao, Y., Stuart, T., Kowalski, M., Choudhary, S., Hoffman, P., Hartman, A., Srivastava, A., Molla, G., Madad, S., Fernandez-Granda, C., et al. (2024). Dictionary learning for integrative, multimodal, and massively scalable single-cell analysis. Nat Biotechnol *42*, 293–304. https://doi.org/10.1038/s41587-023-01767-y.

7. Piñero, J., Bravo, À., Queralt-Rosinach, N., Gutiérrez-Sacristán, A., Deu-Pons, J., Centeno, E., García-García, J., Sanz, F., and Furlong, L.I. (2017). DisGeNET: a comprehensive platform integrating information on human disease-associated genes and variants. Nucleic Acids Res *45*, D833–D839. https://doi.org/10.1093/nar/gkw943.

8. Subramanian, A., Tamayo, P., Mootha, V.K., Mukherjee, S., Ebert, B.L., Gillette, M.A., Paulovich, A., Pomeroy, S.L., Golub, T.R., Lander, E.S., et al. (2005). Gene set enrichment analysis: A knowledge-based approach for interpreting genome-wide expression profiles. Proc Natl Acad Sci U S A *102*, 15545–15550. https://doi.org/10.1073/pnas.0506580102.

9. Street, K., Risso, D., Fletcher, R.B., Das, D., Ngai, J., Yosef, N., Purdom, E., and Dudoit, S. (2018). Slingshot: cell lineage and pseudotime inference for single-cell transcriptomics. BMC Genomics *19*, 477. https://doi.org/10.1186/s12864-018-4772-0.

10. Morabito, S., Reese, F., Rahimzadeh, N., Miyoshi, E., and Swarup, V. (2023). hdWGCNA identifies co-expression networks in high-dimensional transcriptomics data. Cell Rep Methods *3*, 100498. https://doi.org/10.1016/j.crmeth.2023.100498.

11. Yao, Z., van Velthoven, C.T.J., Kunst, M., Zhang, M., McMillen, D., Lee, C., Jung, W., Goldy, J., Abdelhak, A., Aitken, M., et al. (2023). A high-resolution transcriptomic and spatial atlas of cell types in the whole mouse brain. Nature *624*, 317–332. https://doi.org/10.1038/s41586-023-06812-z.

12. Kang, S.H., Fukaya, M., Yang, J.K., Rothstein, J.D., and Bergles, D.E. (2010). NG2+ CNS glial progenitors remain committed to the oligodendrocyte lineage in postnatal life and following neurodegeneration. Neuron *68*, 668–681. https://doi.org/10.1016/j.neuron.2010.09.009.

13. Wang, X., Zeng, C., Wu, Z., Lu, M., Wang, X., Xiu, Y., Wang, Q., Wang, S., Chen, X., Shen, Y., et al. (2025). Chromatin remodeling factor BAF155 coordinates oligodendroglial-neuronal communications linked to regional myelination and autism-like behavioral deficits in mice. Nat Commun *17*, 1165. https://doi.org/10.1038/s41467-025-67930-y.

14. Ting, J.T., Lee, B.R., Chong, P., Soler-Llavina, G., Cobbs, C., Koch, C., Zeng, H., and Lein, E. (2018). Preparation of Acute Brain Slices Using an Optimized N-Methyl-D-glucamine Protective Recovery Method. J Vis Exp, 53825. https://doi.org/10.3791/53825.

15. Schafer, D.P., Lehrman, E.K., Heller, C.T., and Stevens, B. (2014). An engulfment assay: a protocol to assess interactions between CNS phagocytes and neurons. J Vis Exp, 51482. https://doi.org/10.3791/51482.

### Supplementary Figures

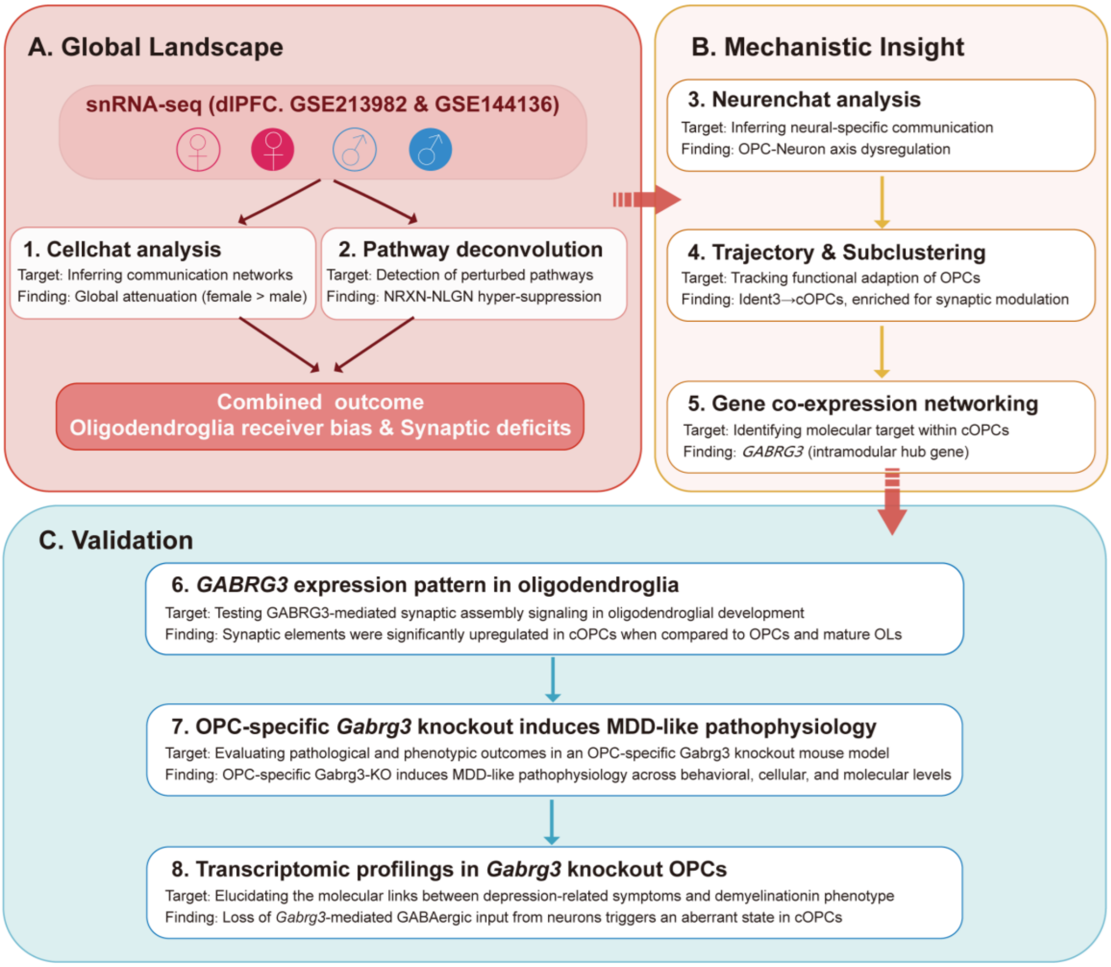


**Supplementary Figure 1. Schematic overview of study design**

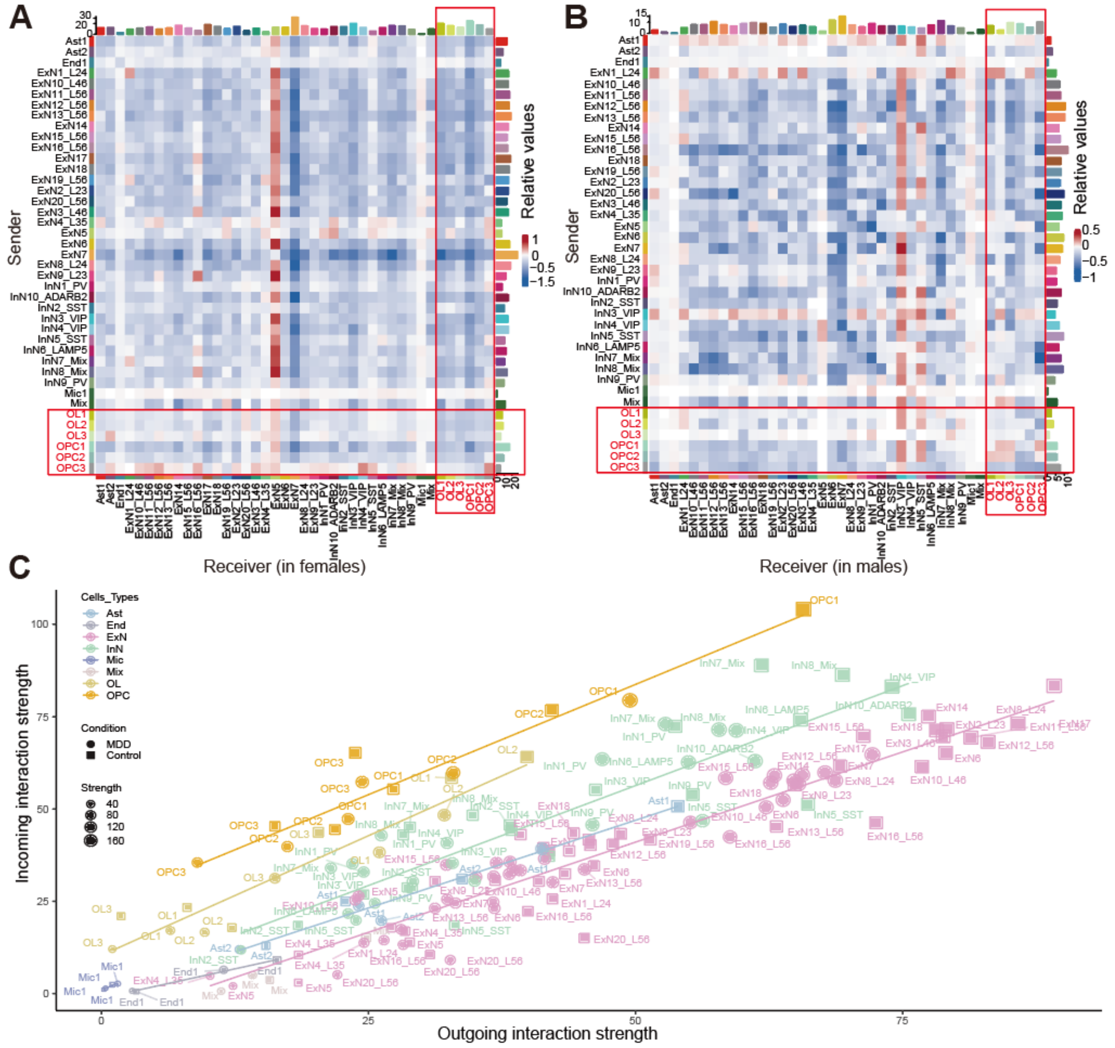


**Supplementary Figure 2. Signatures of cell-type-specific interaction strengths across sexes in MDD**

(A-B) Heatmaps displaying the differential interaction strength among cell clusters between controls and MDD in (A) females and (B) males. Red (or blue) spots represent increased (or decreased) signaling in MDD cohort compared to controls.

(C) Regression mapping of distinct cell clusters within the four cohorts emphasizing the unidirectional receptivity bias of oligodendroglia.

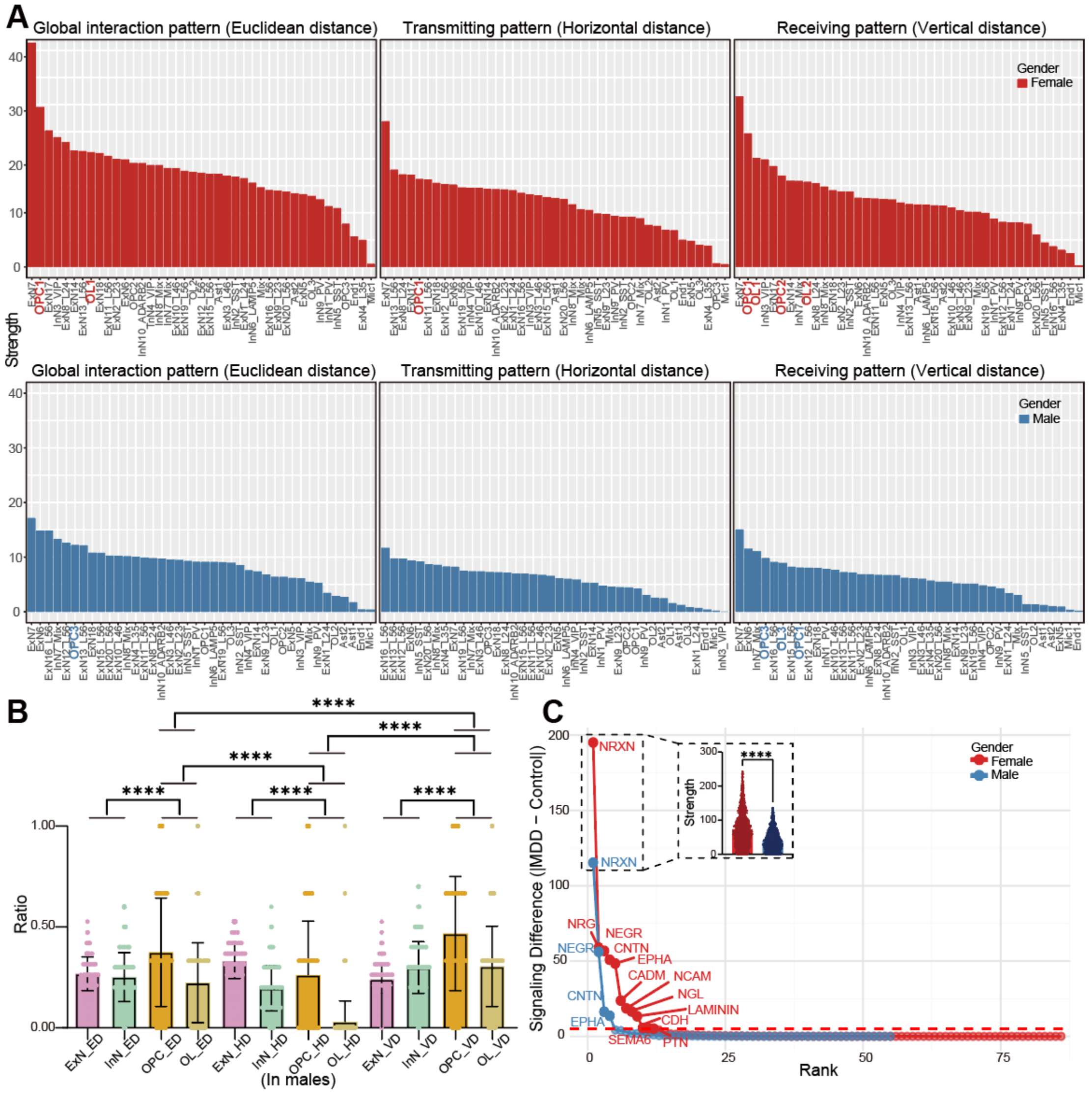

**Supplementary Figure 3. Pathway complexity and interaction divergence profiling across sexes**

(A) Plots quantifying and comparing the interaction divergence between MDD and controls for each cell cluster utilizing Euclidean, horizontal (x-axis), and vertical (y-axis) distances.

(B) Proportional representations of the top 10 most divergent cell clusters in males supporting the distinct functional bias of oligodendroglial lineage cells in signal reception.

(C) Ranking of perturbed signaling pathways by the magnitude of interaction strength changes between MDD and controls in both sexes. The inflection point at a value of 5 (dashed line) serves as the threshold for identifying significant pathway perturbations.

The identification of significantly perturbed signaling pathways was established utilizing the permutation test. For the quantitative comparison, data distributions for (B) and (C) were generated via permutation analysis, and subsequent statistical significance between groups was evaluated using the two-sided Kolmogorov-Smirnov test in GraphPad Prism. *$p < 0.05$, **$p < 0.01$, ***$p < 0.001$, ****$p < 0.0001$.

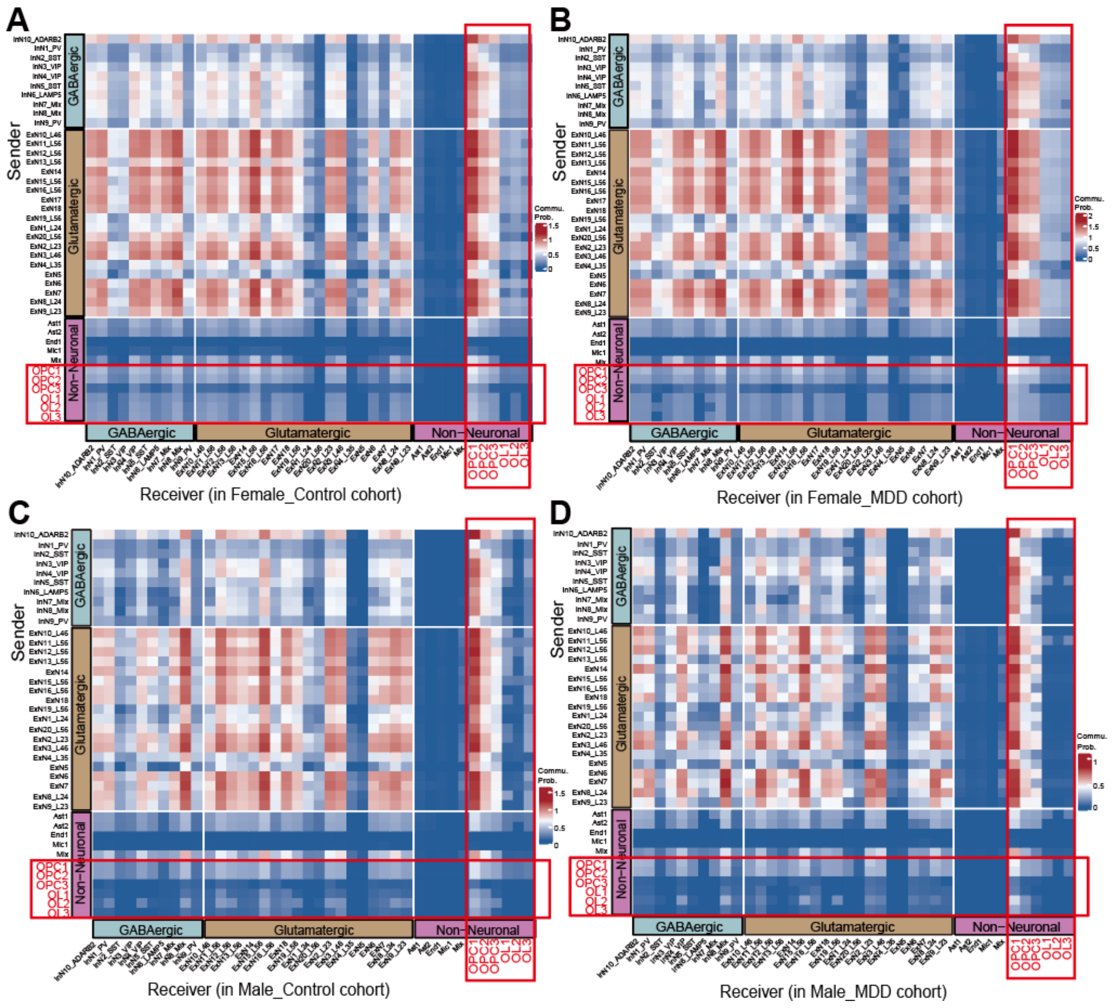


**Supplementary Figure 4. Neural-specific intercellular signaling networks across cohorts**

Heatmaps generated via the NeuronChat framework illustrating the communication probabilities in (A) female controls, (B) female MDD patients, (C) male controls and (D) male MDD patients.

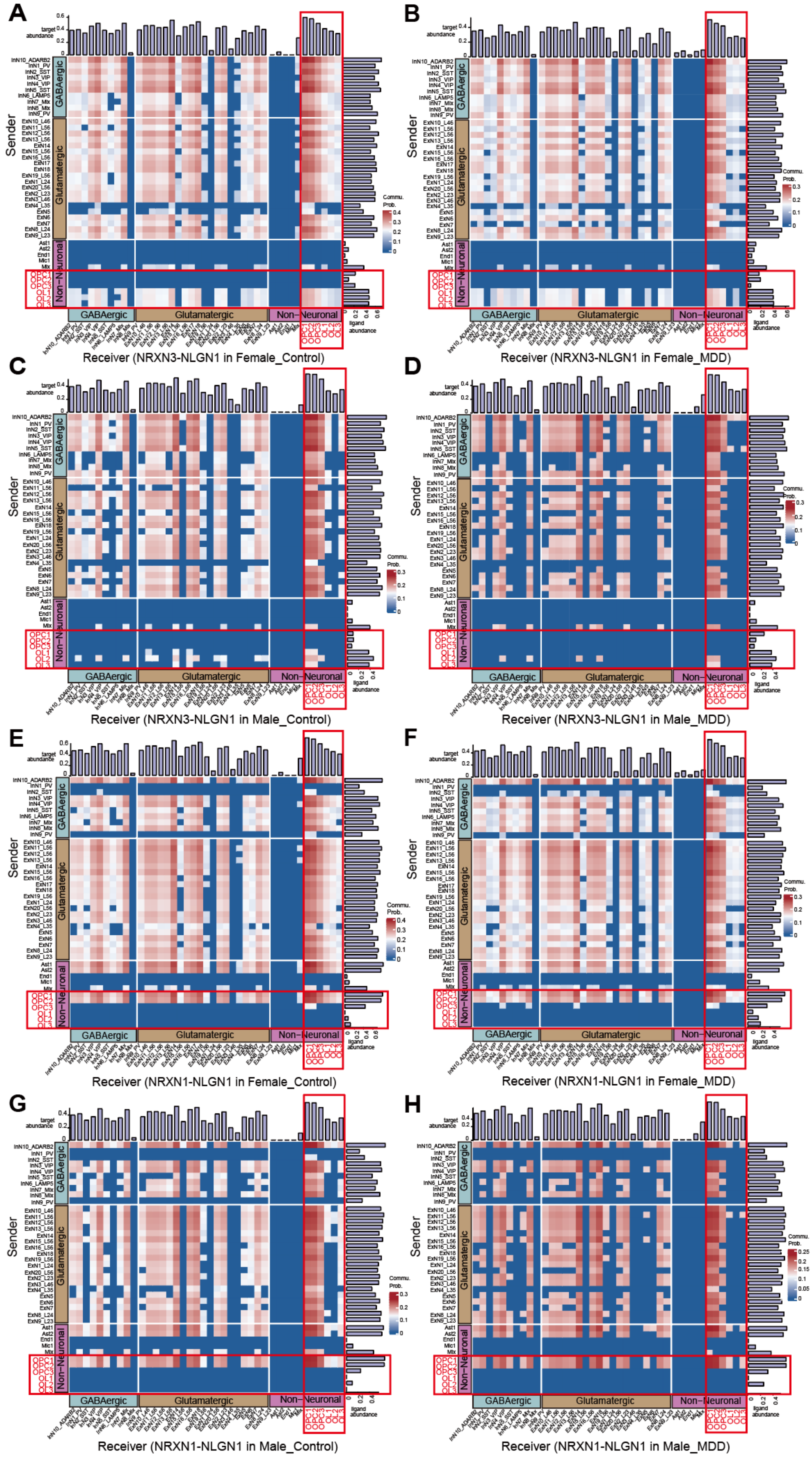
A
B
C
D
E
F
G
H
Sender
Receiver (NRXN3-NLGN1 in Female_Control)
Receiver (NRXN3-NLGN1 in Female_MDD)
Receiver (NRXN3-NLGN1 in Male_Control)
Receiver (NRXN3-NLGN1 in Male_MDD)
Receiver (NRXN1-NLGN1 in Female_Control)
Receiver (NRXN1-NLGN1 in Female_MDD)
Receiver (NRXN1-NLGN1 in Male_Control)
Receiver (NRXN1-NLGN1 in Male_MDD)
GABAergic
Glutamatergic
Non−Neuronal
target abundance
ligand abundance
Commu. Prob.

**Supplementary Figure 5. Interaction mapping of the NRXN-NLGN1 ligand-receptor pairs**

(A-D) Specific intercellular interactions mediated by the single NRXN3-NLGN1 pair within the NRXN pathway in (A) female controls, (B) female MDD patients, (C) male controls and (D) male MDD patients.

(E-H) Specific intercellular interactions mediated by the single NRXN1-NLGN1 pair within the NRXN pathway in (E) female controls, (F) female MDD patients, (G) male controls and (H) male MDD patients.

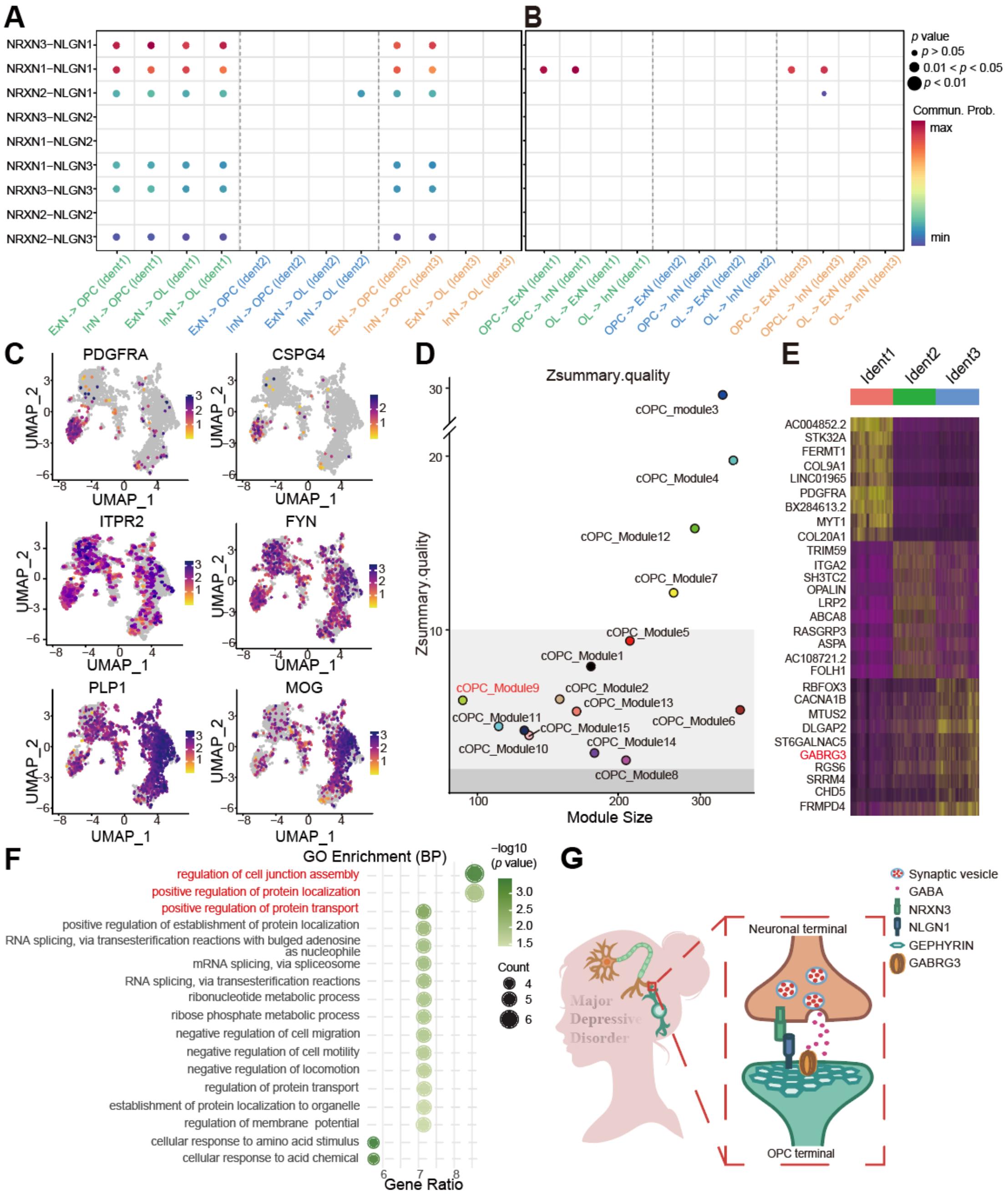


**Supplementary Figure 6. Validation of cOPCs and the *GABRG3*-centered topological subnetwork**

(A) Bubble plot visualizing the communication probabilities of NRXN-NLGN pairs in the neuron-to-OPC interactions.

(B) Bubble plot visualizing the communication probabilities of NRXN-NLGN pairs in the OPC-to-neuron interactions.

(C) Feature plots confirming the cellular identities of oligodendroglial subclusters.

(D) Scatter plot displaying the quality (Z-summary scores) of co-expression modules identified from healthy control cOPCs in MDD.

(E) Heatmap showing the top 10 prioritized transcriptional signatures across oligodendroglial subsets.

(F) GO analysis profiling the biological processes of the enriched genes within the MDD-specific subnetwork of cOPC_Module9.

(G) Diagrammatic summary proposing the possible NRXN3-NLGN-gephyrin-GABRG3 synaptic assembly signaling cascade in neurons.

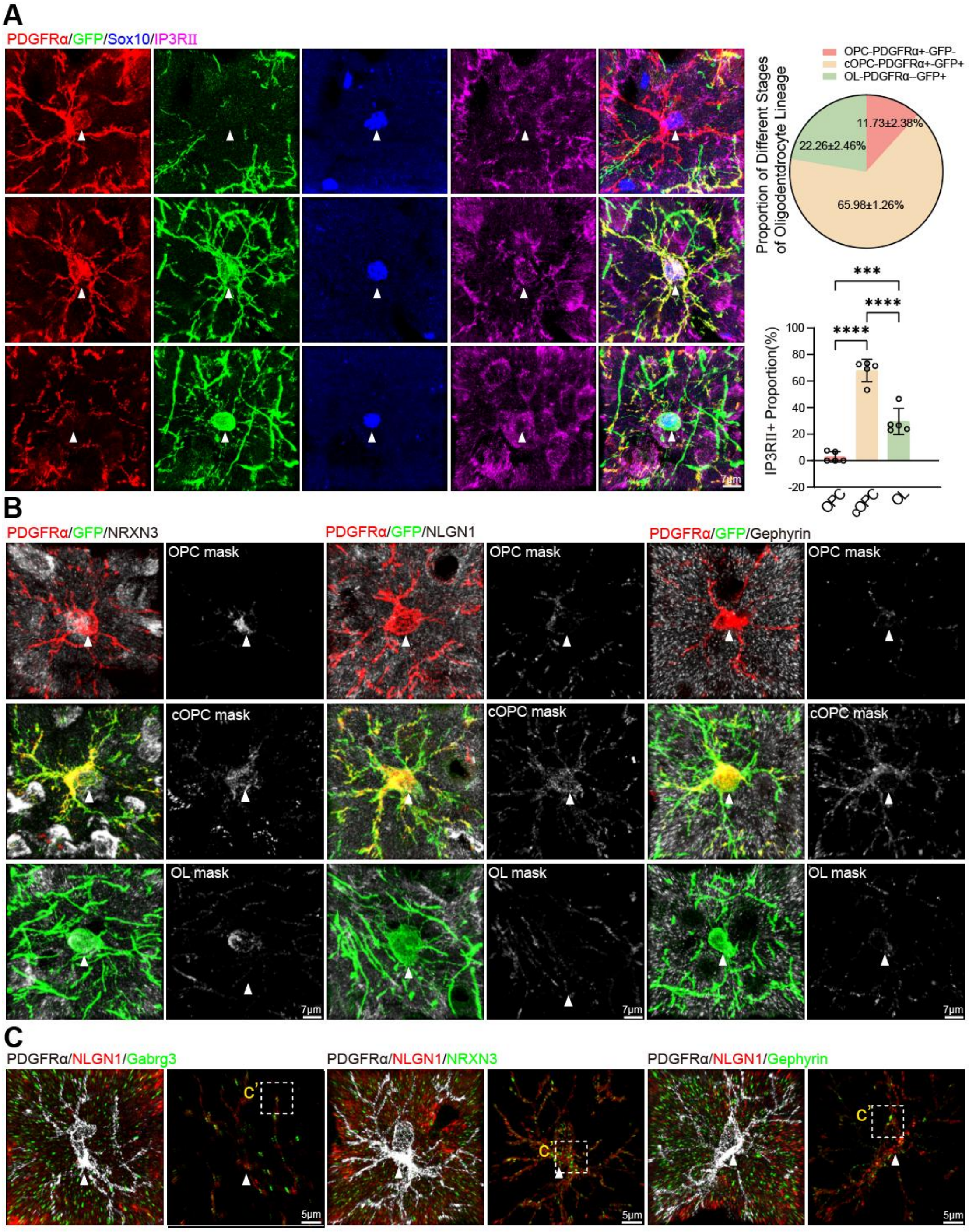


**Supplementary Figure 7. Expression patterns of synaptic elements in oligodendroglia**

(A) Representative images of *NG2*$^{CreERT}$; *Tau-mGFP* mouse brain sections at P14, stained for PDGFRα, IP3R-II, and SOX10. Quantification of GFP signal in PDGFRα- and IP3R-II-positive cells (white arrowheads mark cell bodies). Scale bar = 7 μm, n = 5 mice.

(B) Representative images of PDGFRα, Tau-GFP and synaptic elements (NRXN3, NLGN1 and gephyrin) staining in oligodendroglial lineage cells during their developmental process by super-resolution fluorescence microscopy. The number of synaptic elements per unit of oligodendroglial lineage + area. Scale bar = 7 μm, n=5 mice.

(C) Representative images of synaptic elements in PDGFRα positive areas by super-resolution fluorescence microscopy. Scale bar = 5 μm, n=5 mice.

Data presented as mean ± SEM; ***$p < 0.001$, ****$p < 0.0001$.

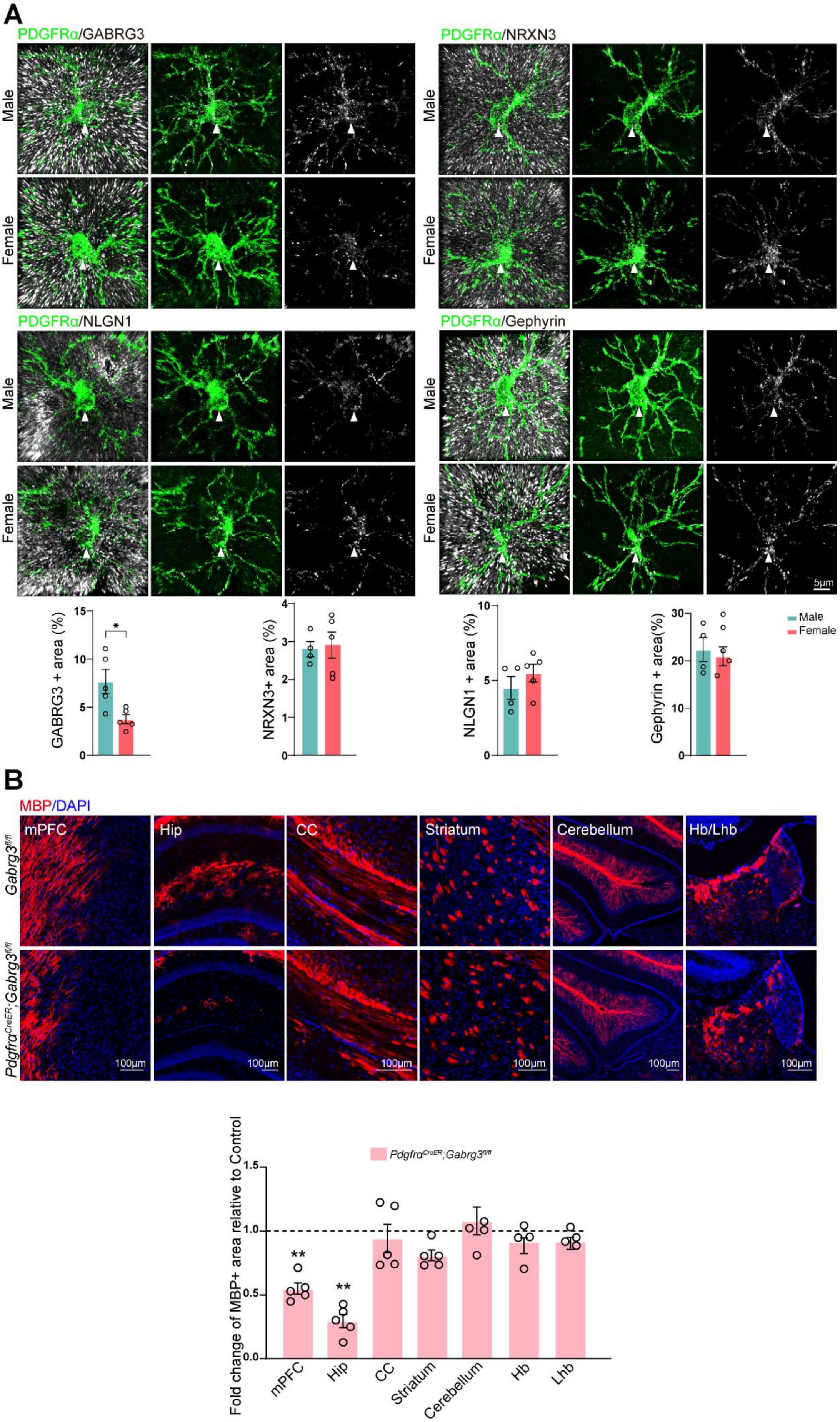
A
PDGFRα/GABRG3
PDGFRα/NRXN3
PDGFRα/NLGN1
PDGFRα/Gephyrin
Male
Female
5μm
GABRG3 + area (%)
NRXN3+ area (%)
NLGN1 + area (%)
Gephyrin + area(%)
Male
Female
B
MBP/DAPI
mPFC
Hip
CC
Striatum
Cerebellum
Hb/Lhb
Gabrg3fl/fl
PdgfraCreER;Gabrg3fl/fl
100μm
Fold change of MBP+ area relative to Control
PdgfraCreER;Gabrg3fl/fl
mPFC
Hip
CC
Striatum
Cerebellum
Hb
Lhb

**Supplementary Figure 8. Sexually divergent synaptic assembly signaling in mice and OPC-specific *Gabrg3* knockout model induce MDD-like pathophysiology**

(A) Representative images of GABRG3, NRXN3, NLGN1 and gephyrin expressions in mice. Quantifications indicate the GABRG3 (also seen in Figure 6F), NRXN3, NLGN1 and gephyrin + area per unit PDGFRα+ area. Scale bars = 5 µm; n = 5 mice.

(B) Representative images and quantifications of MBP staining in the mPFC, striatum, corpus callosum, hippocampus, cerebellum, lateral habenula and habenula of $PDGFR\alpha^{CreER}$; $Gabrg3^{fl/fl}$ and control mice (indicated by dotted line) at P14. Scale bar = 100 µm, n=5 mice.

Data presented as mean ± SEM; $^{**}p < 0.01$.
